# Ablation of endogenously cycling adult cardiomyocytes worsens myocardial function after injury

**DOI:** 10.1101/2020.09.21.306852

**Authors:** Leigh A. Bradley, Alexander Young, Helen O. Billcheck, Matthew J. Wolf

**Author notes:** Corresponding Author: Matthew J. Wolf MD, PhD Associate Professor of Medicine Division of Cardiology University of Virginia Medical Research Building 5 (MR5) Room G213 415 Lane Road Charlottesville, VA 22908.

## Abstract

**Rationale:** Endogenously cycling adult cardiomyocytes (CMs) increase after myocardial infarction (MI) but remain scare, and are generally thought not to contribute to myocardial function. However, this broadly held assumption has not been tested, mainly because of the lack of transgenic reporters that restrict Cre expression to adult CMs that reenter the cell cycle.

**Objective:** We created and validated a new transgenic mouse, *<u>α</u>MHC-Mer<u>D</u>reMer-<u>K</u>i67p-<u>R</u>oxed<u>C</u>re::<u>R</u>ox-<u>L</u>ox-td<u>T</u>omato-e<u>G</u>FP* (denoted *αDKRC*) that restricts Cre expression to cycling adult CMs and uniquely integrates spatial and temporal adult CM cycling events based on the DNA specificities of orthologous Dre- and Cre recombinases. We then created mice that expressed an inducible Diphtheria toxin (DTA), *αDKRC::DTA* mice, in adult cycling CMs and examined the effects of ablating these endogenously cycling CMs on myocardial function after Ischemic-Reperfusion (I/R) MI.

**Methods and Results:** A tandem *αDKRC* transgene was designed, validated in cultured cells, and used to make transgenic mice. The *αDKRC* transgene integrated between *MYH6* and *MYH7* and did not disrupt expression of the surrounding genes. Compared to controls, *αDKRC::RLTG* mice treated with Tamoxifen expressed tdTomato+ in CMs with rare Bromodeoxyuridine (BrdU)^+^, eGFP^+^ CMs, consistent with reentry of the cell cycle. We then pre-treated *αDKRC::RLTG* mice with Tamoxifen to activate the reporter before sham or reperfusion (I/R) myocardial infarction (MI) surgeries. Compared to Sham surgery, the I/R MI group had increased single and paired eGFP+ CMs predominantly in the border zones (5.8 ± 0.5 vs. 3.3 ± 0.3 CMs per ten-micron section, N = 8 mice, n = 16 sections per mouse), indicative of cycled CMs. The single to paired eGFP+ CM ratio was ∼9 to 1 (5.2 ± 0.4 single vs. 0.6 ± 0.2 paired CMs) in the I/R MI group after MI, suggesting that cycling CMs were more likely to undergo polyploidy than replication. The ablation of endogenously cycling adult CMs in *αDKRC::DTA* mice caused progressive worsening left ventricular chamber size and function after I/R MI, compared to controls.

**Conclusions:** Although scarce, endogenously cycling adult CMs contribute to myocardial function after injury, suggesting that these cells may be physiologically relevant.

## Introduction

Although cardiovascular outcomes have improved, CMs, the muscle cells of the heart, are lost even with successful reperfusion, and this loss contributes to adverse LV remodeling, ischemic cardiomyopathy, heart failure, arrhythmia, and death. Heart failure affects 5.7 million Americans with a projected 46% increase in prevalence by 2030, has a ∼50% mortality rate at five years despite current medication and device-based therapies, and accounts for ∼2 million physician office visits and ∼$30 billion in direct medical costs annually.^1^ FDA approved drugs to treat heart failure fall within a few classes designed to dampen the adrenergic or renin-angiotensin signaling systems.^2^ Unfortunately, current therapies can only slow or reverse isolated aspects of heart failure, and there are no reliable therapies available to replace the cardiac muscle loss to MI. Thus, identifying therapeutic targets and drugs to protect myocardium after injury will be groundbreaking, address unmet clinical needs, and represent new strategies to treat cardiovascular diseases.

Mammalian CMs exit the cell cycle during postnatal heart growth.^3, 4^ The total number of CMs in the heart decreases with age, by apoptotic or necrotic cell death, and leads to replacement hypertrophy of the remaining CMs and deposition of extracellular matrix.^5^ CMs experience a variety of insults, including oxidative stress^6^, DNA damage^7^, mitophagy^8, 9^, autophagy^10^, and the effects of inflammation during aging^11^. Protecting CMs from these damaging conditions is one approach to improve or potentially reverse myocardial dysfunction.

Alternatively, enhancing endogenous cycling CMs is another attractive strategy to improve age- and injury-related changes in myocardial function. However, there are several challenges. First, adult cycling CMs are scarce. Adult mammalian CMs undergo limited cell cycling, and even lower rates of proliferation, in response to injury.^12–15^ Human CM turnover is ∼0.04% in the first year of life and ∼0.01% per year in adulthood based on estimates of nuclear bomb test-derived radioisotope decay.^3, 16–18^ CM cycling increases to ∼0.1% in response to injury; however, the turnover is likely overestimated because of polyploidy instead of bona fide replication of CMs.^19^ Second, adult CMs are terminally differentiated and need to integrate electrically, mechanically, and vascularly with surrounding CMs, to repair injured myocardium. Despite these limitations, global enhancement of CM proliferation has been shown to improve myocardial function after injury. For example, Cyclin-D^20, 21^, Hippo/Yap^22–24^, p38^25^, or miRNAs^26^ induced CM proliferation and improved myocardial function after MI. In some cases, as with glycogen synthase kinase (GSK)-3α and GSK-3β knockout mice, enhanced CM cycling causes mitotic catastrophe, a type of cell death that occurs during mitosis.^27^ Thus, merely increasing CM cycling may not translate into beneficial effects.

Accurately distinguishing between proliferation and endoreplication (polyploidy) is required to investigate CM cycling and myocardial function.^28^ Generally, two approaches characterize cycling cells (Supplemental Figure 1). The first provides a temporal “snapshot” of cycling events based on the co-localization of markers expressed in cycling cells, including phospho-histone-3, Ki67, or Aurora B. These markers are expressed only when a cell is actively cycling. Therefore, the detection is dependent on the time a tissue is analyzed after injury and can lead to an over- or underestimation of cycling events. Markers that are specific to the mid-body of cytokinesis may identify proliferative events. The second approach provides a “summation” measurement of cycling events. Typically this approach measures the incorporation of thymidine analogs (BrdU or EdU) into DNA during S-phase using antibodies or Click-It chemistry to identify cycling cells, but cannot distinguish between proliferation and endoreplication (polyploidy). Although combining these two conventional approaches is well-accepted, especially in analyses of large animals and human tissues, there is still an incomplete picture of cell cycling and polyploidy. Clearly, additional approaches are needed to address this fundamental biology.

Transgenic strategies complement detection of markers of cycling and incorporation of thymidine analogs to provide a stronger interpretation of cycling adult CMs, and a few can infer proliferation from polyploidy.^15, 20^ However, new molecular approaches are necessary to drive transgenes specifically in endogenously cycling CMs and interrogate how this scare population of cells may contribute, if at all, to myocardial function. Our overall objective was to create a transgenic mouse that restricted Cre expression to endogenously cycling adult CMs. Our goals were to create a transgenic reporter mouse that fulfilled the following requirements: (1) expression restricted to cycling CMs after exit from the normal developmental cell cycle, (2) adaptability to any LoxP transgenic mouse, (3) a single tandem transgene to facilitate breeding, and (4) avoidance of reporter insertion into the *Rosa26* locus because such an event limits the ability to use the multitude of existing transgenics previously knocked into the *Rosa26* locus, such as RLTG.

To address these challenges, we developed a transgenic approach that utilized the distinct specificities of Dre and Cre recombinases that recognized different DNA recombination sites, sequentially. Dre recombinase is a Type I topoisomerase identified from a screen of P1-like bacteriophage and recognizes “Rox” DNA sites for recombination, similar to, but with different specificity of, Cre recombinase’s recognition of LoxP DNA sites.^29, 30^ Dual Dre and Cre recombinases have been used to lineage trace cell fates, including the contributions of non-myocytes to myocytes, hepatic progenitor cells in liver development, and astrocytes in neurogenesis.^31–35^ Instead of using promoters to label cell lineages, we adapted the technology and created a new transgenic mouse *<u>α</u>MHC-Mer<u>D</u>reMer-<u>K</u>i67p-<u>R</u>oxed<u>C</u>re::<u>R</u>ox-<u>L</u>ox-td<u>T</u>omato-e<u>G</u>FP* (denoted *αDKRC::RLTG*) to identify adult CMs that re-entered the cell cycle based on activation of a Ki67 promoter that drives Cre recombinase. The power of the approach is that a limited exposure to Tamoxifen primes the reporter, allowing experiments to be conducted weeks after recovery from Tamoxifen. CMs re-entering the cell cycle will express Cre at any time after Tamoxifen exposure, and label proliferation or endoreplicating (polyploid) CMs. We observed increases in cycling CMs two weeks after 60 minutes of LAD ligation ischemia followed by reperfusion myocardial infarction (I/R MI) with a single to paired eGFP+ CM ratio was ∼9 to 1 in the I/R MI group after MI, suggesting that cycling CMs were more likely to undergo polyploidy than replication.

Since Cre expression is restricted to CMs that reentered the cell cycle, the *αDKRC* mouse has the potential to drive gene expression in the subpopulation of cycling CMs to investigate signaling pathways that may lead to improved myocardial function during aging or after injury. Endogenous cycling adult CMs after injury are scare and generally thought to not contribute to myocardial function; however, this assumption has not been investigated. Therefore, we used *αDKRC* mice induce Diphtheria toxin (*αDKRC::DTA*) and ablate CMs that reentered the cell cycle after I/R MI and observed worsened LV systolic function in *αDKRC::DTA* that were pretreated with Tamoxifen, compared to controls. Collectively, *αDKRC* represents the next evolution in Ki67 promoter-driven Cre recombinases^36, 37^ and provides opportunities to characterize the subpopulations of endogenously cycling adult CMs.

## Methods

### Plasmids

αMHC-mCherry-Rex-Blasticidin (Addgene plasmid #21228; http://n2t.net/addgene:21228; RRID: Addgene_21228) was a gift from Mark Mercola.^38^ pCAG-NLS-HA-Dre (Addgene plasmid #51272; http://n2t.net/addgene:51272; RRID: Addgene_51272), pCAG-roxSTOProx-ZsGreen (Addgene plasmid #51274; http://n2t.net/addgene:51274; RRID: Addgene_51274) and pCAG-Roxed-Cre (Addgene plasmid #51273; http://n2t.net/addgene:51273 ; RRID:Addgene_51273) were gifts from Pawel Pelczar.^32^ pLV-CMV-LoxP-DsRed-LoxP-eGFP (Addgene plasmid #65726; http://n2t.net/addgene:65726 ; RRID: Addgene_65726) was a gift from Jacco van Rheenen.^39^ pCAG-Cre (Addgene plasmid #13775; http://n2t.net/addgene:13775 ; RRID:Addgene_13775) was a gift from Connie Cepko.^40^ pSF-CAG-Kan (OG505) CAG promoter vector (Oxford Genetics, Inc). pCAGG-Fucci2a plasmid was obtained from Riken (RDB13080).^41^

### Generation of pCAG-MerDreMer, pCAG-Rox-STOP-Rox-LoxP-DsRed-Stop-LoxP-eGFP-Stop, Ki67-Fucci, and αMHCp-MerDreMer-Ki67p-RoxedCre (αDKRC) plasmids

#### pCAG-MerDreMer

A new multi-cloning site was created (MCS SC40pA) (see list of DNA tiles in Detailed Methods) and subcloned into pSF-CAG-Kan (OG505) CAG promoter vector (Oxford Genetics, Inc) using BglII and NheI restriction enzymes to remove the CAG promoter and multi cloning site (MCS) thereby generating an OG-MCS Sv40pA plasmid with new MCS functionality. G525R mutant forms of the mouse estrogen receptor (Ert1 and Ert2) and hemagglutinin (HA) epitope-tagged Dre recombinase (HA-Dre) tiles were synthesized by Gibson DNA assembly (IDT DNA), confirmed by Sanger DNA sequencing, and sequentially subcloned into OG-MCS Sv40pA plasmid to make the plasmid encoding a tamoxifen-inducible Dre (denoted MerDreMer). The HA-Dre DNA tile was designed based on pCAG-NLS-HA-Dre (Addgene Plasmid #51272).^32^ The CAG promoter was subcloned from pSF-CAG-Kan (OG505) CAG promoter vector into the MerDreMer plasmid to generate pCAG-MerDreMer (Supplemental Figure 2).

#### pCAG-Rox-STOP-Rox-LoxP-DsRed-Stop-LoxP-eGFP-Stop

Briefly, pSF-CAG-Kan (OG505) CAG promoter vector (Oxford Genetics, Inc) was digested with HindIII and BamHI to create a new plasmid containing a HindIII-SalI-NcoI-BstBI-AgeI-BamHI MCS (designated CAG-HindIII-SalI-NcoI-BstBI-AgeI-BamHI plasmid). The Rox-STOP-ROX cassette was removed from CAG-Rox-STOP-Rox-zfGreen using SalI and NcoI restriction enzymes and subcloned into the CAG-HindIII-SalI-NcoI-BstBI-AgeI-BamHI plasmid. Then, the LoxP-dsRED-LoxP-eGFP cassette was removed from pLV-CMV-LoxP-DsRed-LoxP-eGFP using BstBi and AgeI restriction enzymes and subcloned into the CAG-HindIII-SalI-NcoI-BstBI-AgeI-BamHI plasmid containing the Rox-STOP-ROX cassette. The resultant pCAG-Rox-STOP-Rox-LoxP-DsRed-Stop-LoxP-eGFP-Stop plasmid was validated by Sanger DNA sequencing (Supplemental Figure 2).

#### Ki67p-Fucci

The 1,524 bp Ki67 promoter described by Zambon^42^ was isolated from *C57B6J* genomic DNA by PCR, modified to produce 5P BglII-NheI and 3-HindII restriction sites, validated by Sanger DNA sequencing, and subcloned into OG-MCS-SV40pA. Next, Fucci2aR was subcloned into the Ki67p-OG-MCS-SV40pA using EcoRI and KpnI restriction sites and validated by DNA Sanger sequencing (Supplemental Figure 2).

#### pCAG-MerDreMer-Ki67p-RoxedCre and αMHCp-MerDreMer-Ki67p-RoxedCre (*αDKRC*)

The mouse αMHC promoter was isolated from αMHC-mCherry-Rex-Blasticidin using 5P and 3P primers continuing SalI restriction sites, sequenced, and subcloned into the plasmid encoding MerDreMer to generate the plasmid encoding αMHCp-MerDreMer. The plasmid encoding Ki67p-Rox-STOP-Rox-Cre was generated as follows. The 1,524 bp Ki67 promoter was isolated from *C57B6J* genomic DNA by PCR, modified to produce 5P BglII-NheI and 3-HindII restriction sites, validated by Sanger DNA sequencing, and subcloned into OG-MCS-SV40pA. The Rox-Stop-Rox cassette was subcloned from pCAG-Roxed-Cre using primers containing 5P EcoRI and 3P KpnI restriction sites, validated by Sanger DNA sequencing, and subcloned into the Ki67p-OG-MCS-SV40pA plasmid (designated “Ki67p-RoxedCre” plasmid). Ki67p-RoxedCre cassette was subcloned from the Ki67p-RoxedCre plasmid into the αMHCp-MerDreMer plasmid using NheI restriction enzyme to generate αMHCp-MerDreMer-Ki67p-RoxedCre plasmid. The αMHCp-MerDreMer-Ki67p-RoxedCre plasmid was validated by Sanger DNA sequencing. CAG-MerDreMer-Ki67p-RoxedCre was generated in similarly except the CAG promoter from the pSF-CAG-Kan (OG505) vector was used instead of αMHC promoter (Supplemental Figure 2).

#### Cell Culture

HEK293T cells were cultured in DMEM supplemented with 10% fetal bovine serum. For experiments testing the constructs encoding Dre and Cre recombinases, HEK293T cells were transfected with 1 ug of pCAG-Rox-STOP-Rox-LoxP-DsRed-Stop-LoxP-eGFP-Stop and indicated plasmids using Lipofectamine 2000 (Invitrogen, Inc.). 24 hours after transfections, cells were treated with 2 uM of Tamoxifen or PBS vehicle control for an additional 24 hours and dsRed and eGFP were directly visualized using a Leica DM2500 fluorescence microscopy system. For experiments testing the mouse Ki67 promoter, HEK293T cells were transfected with Ki67p-fucci, cultured for 48 hours, and fixed in 4% paraformaldehyde. Immunofluorescent staining was performed using anti-mCherry, anti-mVenus, and anti-Ki67 antibodies (see Table in Detailed Methods). Fucci and anti-Ki67 antibody signals were co-localized and quantified using ImageJ.

#### Generation of transgenic mice harboring *αDKRC*

PvuI and SwaI restriction enzymes were used to generate the 13,480 bp linear DNA fragment continuing αDKRC for *C57BL/6J* pronuclear plasmid injection to generate transgenic founder mice by Cyagen, Inc. After multiple rounds of injections, three founder mice were identified as harboring the αDKRC transgene (Figure 1).

**Figure 1.**
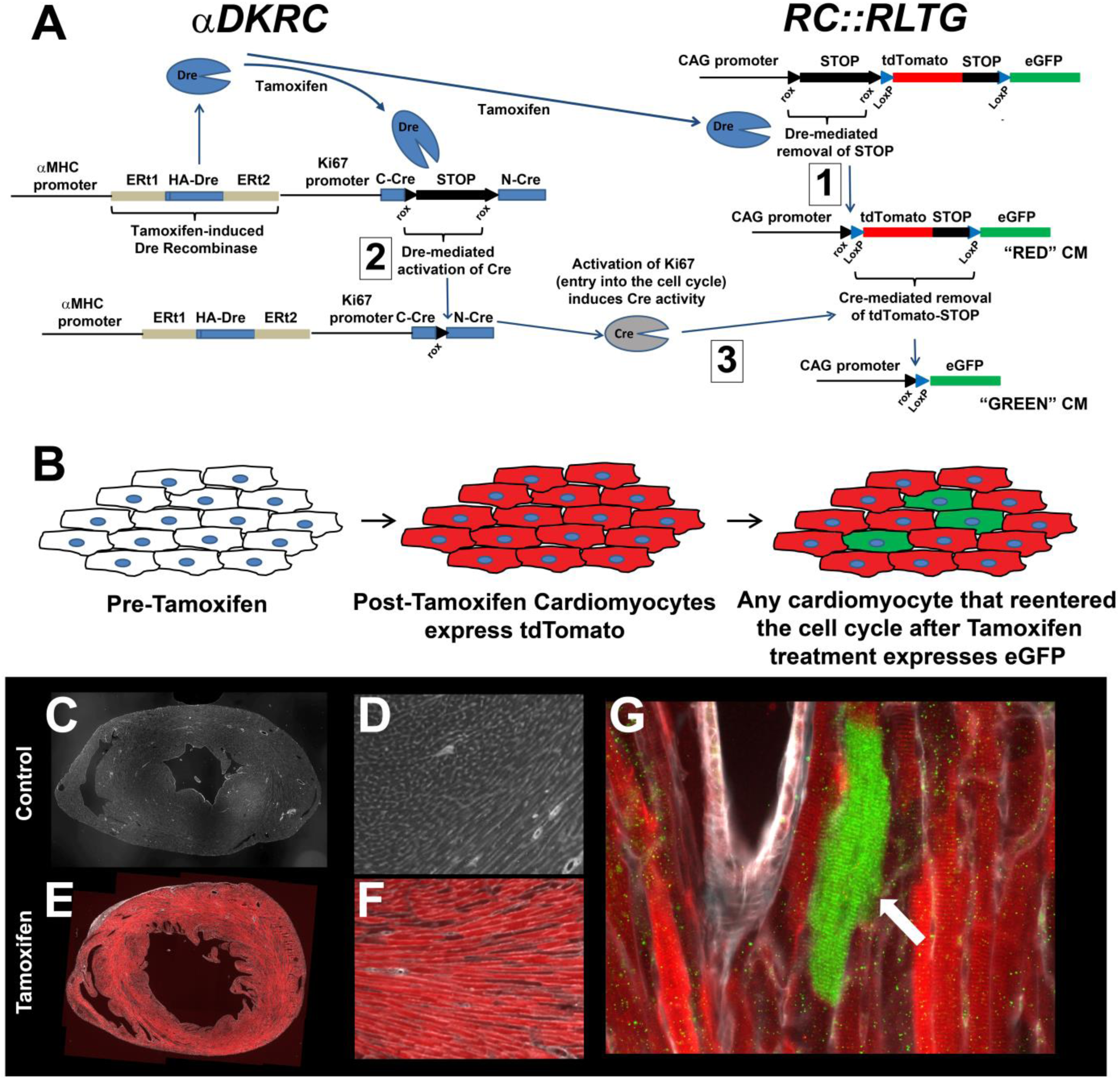
*αDKRC::RLTG* transgenic mice. (A and B) Overview of the labeling of cycling adult CMs using *αDKRC::RLTG* transgenic mice. Tamoxifen exposure induces the CM-specific of Dre-recombinase and subsequent excision of STOP cassettes from the RLTG reporter (Box labeled 1), resulting in the expression of tdTomato and RoxedCre (Box labeled 2) to generate catalytically active Cre recombinase under control to the Ki67 cell cycle promoter. CMs reentering the cell cycle, as defined by activation of the Ki67 promoter, express Cre (Box labeled 3) and excise the tdTomato-STOP cassette resulting in the expression of eGFP in CMs. A short duration of Tamoxifen exposure activates that reporter system. Schematic of expected changes in tdTomato and eGFP labeled CMs. **(C-F)** Representative immunofluorescence images of a heart from *αDKRC::RLTG* transgenic mice treated with peanut oil (C & D) or Tamoxifen (1 mg/kg IP daily x 5 d) (E & F). Paraffin sections were stained for tdTomato (Red), eGFP (Green), and WGA (Gray). **(G)** Representative immunofluorescence image from panel E with the white arrow pointing to a single eGFP^+^ CM identified by the review of eight 10-micron short-axis heart sections. Paraffin sections were stained for tdTomato (Red), eGFP (Green), and WGA (Gray).

Genotyping was performed using PCR across four regions of the transgene using the following primer sets: Transgene PCR primer F1: AGAGCCATAGGCTACGGTGTA; Transgene PCR primer R1: ATGGAGGGTCAAATCCACAAAGC; Annealing Temp: 60^0^C; Transgene PCR product size: 419 bp. Transgene PCR primer F2: AGCATGAAGTGCAAGAACGTG; Transgene PCR primer R2: GGTGAAGTGAATGGAGCCTAAAC; Annealing Temp: 60^0^C; Transgene PCR product size: 444 bp. Transgene PCR primer F3: CGAGCGGTGGTTCGACAAGTG; Transgene PCR primer R3: AACCTCTACAAATGTGGTATGGCTG; Annealing Temp: 60^0^C; Transgene PCR product size: 354 bp. Transgene PCR primer F4: CGCATTGTCTGAGTAGGTGTCATTC; Transgene PCR primer R4: CACTGCATTCTAGTTGTGGTTTGTC; Annealing Temp: 60^0^C; Transgene PCR product size: 372 bp.

Of the three *αDKRC* founders, one failed to produce genotype positive offspring. One produced genotype positive offspring but did not produce transgenic protein. The third produced genotype positive offspring, produced cardiomyocyte-specific transgenic Dre after tamoxifen exposure, and was bred into *C57BL/6J* genetic background for all subsequent experiments.

#### Mapping the αDKRC transgene insertion

The αDKRC transgene insertion site was mapped using nested inverse PCR and Sanger DNA sequencing. Briefly, *C57BL/6J* and *αDKRC* genomic DNA was isolated (DNAeasy, Qiagen Inc.), digested with CviQI restriction enzyme, and ligated with T4 ligase to concatemerize DNA fragments. Nested PCR was performed using the following primer sets:

First round PCR for 5P end:

5P5PRev1 TGTGTGTAACGCAACGATTGATAG
5P3PFor1 CACATCCATCTTCGATGGATAGCG.

Second round PCR for 5P end:

5P5PRev2 GGGCCTCTAAACGGGTCTTGAG’
5P3PFor2 TTATCTAACTGCTGATCGAGTGTAG

First round PCR for 3P end:

3P5Prev1 TTATGCTACGATACCGATAGAGATG
3P3Pfor1 CCTTAAGGTCGTCAGCTATCCTG.

Second round PCR for 3P end:

3P5Prev2 CCACATTTGTAGAGGTTTTACTTG
3P3Pfor2 TCGATCAAGACATTCCTTTAATGG

Unique PCR products obtained from αDKRC genomic DNA were purified after agarose gel electrophoresis, subcloned into pCR-Blunt II-Topo vectors (Invitrogen, Inc.), and analyzed by Sanger sequencing. A single insertion site was identified on Chromosome 14 between bp 54,967,899 and 54,969,800 (Mouse Genome GRCm38, Ensembl Genome Browser) with the corresponding 1,901 bp of genomic DNA deleted.

#### Quantitative PCR (qPCR)

Cardiomyocytes from eight-week old *αDKRC/+* and control genotype negative littermates (5 hearts per group) were isolated by retrograde perfusion of collagenase using a Langendorff apparatus.^43^ RNA was extracted from the heart tissue using the Rneasy kit from Qiagen. For reverse transcription, random primers (1μg) and 10 ng of total RNA were used in a final reaction volume of 20μl containing 100 units of Superscript II (Invitrogen). PCR was performed in duplicate for 40 cycles using 10% of the volume of the reverse transcription in a total volume of 25μl that included 12.5μl of Taqman Gene Expression Master Mix (4369016 Applied Biosystems) and a 1x final concentration of the following Taqman Gene Expression Assays (Applied Biosystems) for Myh6 (Mm00440359_m1), Myh7 (Mm00600555_m1), Mhrt2 (Mm01320179_g1) were obtained from ThermoFisher Scientific. mRNA was quantified by real-time PCR analysis using the CFX96 Real Time System coupled with the C1000 Touch Thermal Cycler (BioRad, Inc.). The ΔCT method was used to quantify all relative mRNA levels using 18S RNA as the reference and internal standard. The TaqMan primer-probe set for 18S RNA (4310893e) with the Vic/Tamra detection system was used to measure 18S RNA in replicate samples to those taken for the above listed mRNA quantifications.

#### Mice

*C57BL/6J* (JAX stock #000664), *RC::RLTG* (*B6.Cg-Gt(ROSA)26Sor^tm1.2^(CAG–tdTomato,–EGFP)^Pjen^/J*)^34^, *αMHC-Cre (B6.FVB-Tg(Myh6-cre)2182Mds/J)*^44^, and ROSA-DTA (*B6.129P2-Gt(ROSA)26Sor^tm1(DTA)Lky^/J*)^45^ mice were purchased from Jackson labs. *CAG-STOP-Fucci2aR* was obtained from the European Mouse Mutant Archive. *CAG-STOP-Fucci2aR* and *αMHC-Cre* were backcrossed nine generations into *C57BL/6J* before establishing *αMHC-Cre/+::CAG-STOP-Fucci2aR* mice. All mice were housed and maintained in accordance with UVA Animal Care and Use Committee approved protocols (UVA Animal Care and Use Committee (ACUC) Wolf Lab protocol #4080).

To test for αDKRC expression, eight to ten week old male and female mice harboring the αDKRC transgene were treated with peanut oil vehicle or Tamoxifen (1 mg/kg) IP daily for 5 days. Hearts were removed, fixed in 10% Neutral Buffered Formalin (NBF) prior to embedding in paraffin. Ten micron short axis sections were prepared and stained with anti-HA, anti-cardiac Troponin, and wheat germ agglutinin (WGA) to identify cardiomyocyte specific, tamoxifen-dependent expression of recombinant MerDreMer protein.

Eight to ten week old male and female *αDKRC/+; RC::RLTG/ RC::RLTG* mice were used for experiments. Mice were treated with peanut oil vehicle or Tamoxifen (1 mg/kg) IP daily for two five-day periods with a two-day rest between rounds (i.e., five days “on”, two days “off”, and five days “on”) followed by a two-week recovery period prior to indicated experiments.

#### Left Anterior Descending (LAD) I/R MI surgery

All animal care and surgeries were in accordance with UVA ACUC Policy on Rodent Surgery and Perioperative Care under ACUC-approved animal protocol (UVA ACUC Wolf Lab protocol #4080). Procedure: Male and female twelve to sixteen week old mice were weighed and then anesthetized in an induction chamber using the gas anesthetic Isoflurane (3% volume/weight, oxygen 500ml/min). The animal was placed in the supine position on a face mask connected to the anesthesia system. Isoflurane was adjusted to provide a maintenance level (1.8-2.2% volume/weight, oxygen 500ml/min) throughout the procedure. Anesthesia was monitored closely by the following methods: (1) Depth and rate of respiration; (2) Heart rate by ECG; (3) Mucous membrane color; (4) Body Temperature via Physio Suite Homoeothermic Temperature Monitoring System; (5) Reflexes & toe pinch; (6) Overall appearance of muscle relaxation. All surgical procedures were carried out with a stereo microscope exclusively for small animal surgery. Normal body temperature was be maintained using an electrical heating pad on a feedback system via rectal probe.

To prepare for surgery, the mouse was first given subcutaneous fluids (veterinary Normosol), and Atropine as a pre-anesthetic to decrease mucous secretion and to prevent gag reflex during intubation. The neck and chest wall was shaved, and then prepped three times with alternating wipes of povidone-iodine and 70% ethanol. Mice were placed in the supine position on a heating pad and an endotracheal intubation was performed under direct laryngoscopy with a lighted fiber optic stylus to visualize the vocal cords and insert tracheal tube. The endotracheal tube was then connected to the automated ventilator (Kent Scientific, Inc.) and the mouse was ventilated (tidal volume = 1.0 mL, rate = 120 breaths/minute). Bupivacaine was infiltrated subcutaneously into the area of the left 3^rd^ and 4^th^ intercostal space. A left anterior thoracotomy was performed using sterile technique by making a small subcutaneous incision lateral from the sternum with sharp scissors. The pectoralis muscle groups were identified and separated by blunt dissection, and held open using fixed retractors. The thoracic cage was exposed and the 3^rd^ intercostal space was identified. Incision into the 3^rd^ intercostal space was done by a combination of blunt micro-scissors and micro-cauterizing tool in order to minimize bleeding. Retractors were then repositioned onto the upper and lower ribs in order to visualize the heart. The pericardium was then blunt dissected in order to expose the anterior view of the heart. The LAD on the surface of the left ventricular wall was identified and an 8-0 prolene suture was placed through the myocardium into the anterolateral LV wall underneath the LAD at the level of the lower atrium (1 mm below the left auricle), and the suture was tied. Occlusion of blood flow is observed by blanching of the heart muscle and ST elevation confirmed by ECG. After 60 minutes the ligature was removed. The chest wall, facial planes, and skin were sutured and the mouse was recovered and treated with analgesics. In sham control mice, the entire procedure was identical except for the ligation of the LAD.

#### Isolation of CMs

CMs from twelve-week old *αDKRC/+* and littermate controls were isolated by retro-aortic cannulation and collagenase treatment by Langendorff isolated heart perfusion as described by Simpson.^43^

#### BrdU and EdU Administration via osmotic Minipump

BrdU Sigma (B5002) solution was prepared aseptically in a sterile vial the day of the procedure, by dissolving 25 mg of BrdU in 1mL of sterile USP grade DMSO (Sigma #D1435) and water. The pH of the solution was in the physiological range. The BrdU solution was filtered using a 0.22 micron filter prior to loading into sterile, pre-packaged Azlet mini-pumps (#2004) that was subcutaneously implanted into mice. For Experiments using EdU, a 10 mg/ml stock of EDU (Sigma #900584) was prepared in water:DMSO solution (50:50 by volume), filtered and loaded into pre-packaged Azlet mini-pumps (#2004).

#### Histology and Immunohistochemistry

Hearts were excised and fixed in 10% Neutral Buffered Formalin (Fisher, Inc.) for a minimum of four hours prior to embedding in paraffin. Ten micron sections were prepared in short axis orientation by microtome with 8 sections per glass slide. Paraffin was removed and the tissue sections were rehydrated using Xylene and serial ethanol wash steps, respectively. Antigen retrieval was performed by incubating tissue sections in boiling 1x Unmasking solution (Vector Labs H-3300) for 22 minutes. After cooling to room temperature, the tissue sections were treated with Sudan Black to quench auto-fluorescence. Briefly, tissue sections were incubated in 0.1% Sudan Black (Sigma, Cat# 199664) in 1x PBS and 70% ethanol at room temperature for 20 minutes to quench auto fluorescence, followed by 3 x five minute washes in 1x PBS containing 0.02% Tween 20, and a final five minute wash in 1x PBS.^46^

For immunofluorescence staining, tissue sections were blocked in 1X PBS containing fish skin gelatin oil (FSG) + Donkey Serum (serum : buffer, 1:10) for 1 hour at room temperature. For primary antibodies derived from mice, an additional step to block endogenous mouse IgG was performed using 1X PBS containing Goat F(ab) Anti-Mouse IgG H&L (Abcam-ab6668) for 1 hour at room temperature. After washing 3 x 2min in 1x PBS containing FSG and 0.1 %Tween-20, tissue sections were incubated in primary antibodies (see table for dilutions) in 1x PBS contacting FSG overnight at 4°C. Tissue sections were washed with 1x PBS containing FSG and 0.1 %Tween-20 for 5 minutes followed by 1X PBS for 5 min prior to incubation with secondary antibodies (see table for dilutions) in 1x PBS containing FSG and 0.1 %Tween-20. Secondary antibodies were removed and sections were washed with 1x PBS containing FSG and 0.1 %Tween-20 for 5 minutes followed by 1X PBS for 5 min prior to imaging. For nuclear staining, DRAQ7 (BD Biosciences 564904) in 1:20 in 1X PBS was used for 5 minutes at room temperature. WGA 647 (Invitrogen W32466) was used at a dilution of 1:100 in 1xPBS for staining cell membranes. EdU was detected using Click-iT™ EdU Cell Proliferation Kit for Imaging, Alexa Fluor™ 594 dye (Invitrogen C10339).

#### Quantification of CM endoreplication and proliferation

Images were obtained using a Leica DM2500 Fluorescence microscopy system with a Leica DFC7000 T fluorescence color camera or a Leica THUNDER imaging system and Leica LAS X Multi Channel Acquisition software. For imaging, fluorescence channels were calibrated to background of control sections stained with secondary antibodies alone. eGFP^+^ CMs visualized using an YFP filter and quantified from 6-8 ten-micron short axis sections of three slides per animal separated by ∼400 microns per slide. The infarct zone was identified by WGA. The non-infarct zone was divided into thirds with the two regions adjacent to the infarct defined as the border zones and the middle region defined as the remote zone (Supplemental Figure 3). eGFP^+^ CMs present in the same location on sequential slides were counted once among all sections to avoid over-representation of eGFP^+^ cells. CM endoreplication was defined as single GFP+ cells, indicative of cell cycle reentry without replication, and proliferation as two neighboring GFP+ CMs separated by cell membranes stained by WGA. Total CMs per section were calculated using ImageJ.

#### Echocardiography

Mice were anesthetized with Isoflurane and echocardiography was performed using a Vevo 1100 (Visual Sonics) system. Images were analyzed using VevoView software.

#### Statistics

GraphPad Prism 8 (GraphPad Software, Inc.) was used for statistical analyses. Student’s t-tests and AVONA with Bonferroni correction for multiple comparisons were used to calculate p-values.

## Results

### Generation and validation of a tandem tamoxifen-inducible Dre and cell cycle-dependent Roxed Cre recombinases

Ki67 is a marker of cell cycle entry.^42, 47, 48^ Therefore, we tested the fidelity of a Ki67 promoter to label cycling cells intending to design a transgenic reporter mouse to label cycling CMs that was readily adaptable to existing loxP-based transgenics. A 1.5 kb mouse Ki67 promoter^42^ was isolated from *C57BL/6J* genomic DNA and used to drive a 2^nd^ generation Fluorescence Ubiquitin Cell Cycle Indication (Fucci2aR)^41^ encoding bicistronic mCherry-hCdt1 (amino acids 1 -130) and mVenus-hGeminin (hGem) (amino acids 10-110) to validate the Ki67 promoter in cultured cells (Supplemental Figures 2). The expression of mVenus and mCherry-derived Fucci2aR driven by the 1.5 kb mouse Ki67 promoter occurred only in HEK293T cells that also expressed endogenous Ki67 protein expression as detected by anti-Ki67 antibodies, suggesting that the 1.5 kb Ki67 promoter was sufficient to label cycling cells (Supplemental Figure 4).

Next, Dre and Cre were engineered to validate the specificities of each enzyme in the presence of Rox and LoxP recognition sites because of the planned intention to use sequential recombinases to label cycling CMs. A CAG-Rox-STOP-Rox-LoxP-dsRed-STOP-LoxP-eGFP-STOP reporter plasmid (denoted Rox/Lox-Red/Green reporter plasmid) was generated to test the recombinases (Supplemental Figures 5). The presence of active Dre-recombinase produces dsRed expression because of the removal of a Rox-STOP-Rox cassette from the Rox/Lox-Red/Green reporter plasmid. The subsequent expression of active Cre-recombinase then excises the dsRed cassette, producing eGFP expression.

Tamoxifen-induced transgene activation of Dre recombinase was next designed and tested, similar to the characterization of Tamoxifen-induced Cre (MerCreMer) strategies. A tamoxifen-inducible Dre recombinase (designated *MerDreMer*) was created using mammalian estrogen receptors (Supplemental Figures 2). A Dre-dependent Cre was generated that encoded a Rox-STOP-Rox cassette between amino acids 59 and 60 of Cre (denoted RoxedCre) and validated using constitutive CAG promoter or the 1.5 kb Ki67 promoter. (Supplemental Figure 2). Recombinant MerDreMer protein required Tamoxifen for catalytic activity and was specific for Rox sites in HEK293T cells co-transfected with the Rox/Lox Red/Green reporter plasmid (Supplemental Figure 6). In the presence of Tamoxifen, HEK-293T cells co-transfected with CAG-MerDreMer, CAG-RoxedCre, and Rox/Lox Red/Green reporter expressed eGFP without dsRed, consistent with the sequential excision of Rox- and LoxP flanked cassettes in the reporter. Additionally, HEK-293T cells co-transfected with CAG-MerDreMer, Ki67p-RoxedCre, and Rox/Lox Red/Green reporter expressed dsRed and eGFP consistent with cell cycle-dependent expression of eGFP.

Based on the results of co-transfections of individual plasmids encoding CAG-MerDreMer and Ki67p-RoxedCre, a tandem CAG-MerDreMer-Ki67p-RoxedCre plasmid was generated (Supplemental Figures 2) and tested in transiently transfected HEK-293T cell using the Rox/Lox- Red/Green reporter (Figure 6A). dsRed and eGFP were not detected in the absence of Tamoxifen (vehicle alone); however, dsRed and eGFP were expressed after Tamoxifen treatment, consistent with activation of MerDreMer and subsequent expression of Ki67p driven, Dre-dependent Cre-recombinase. The presence of double-positive dsRed and eGFP cells was consistent with the half-life of dsRed protein in cells. The results supported the Tamoxifen-inducible activation of Dre-recombinase and cell cycle-dependent activation of Cre driven using the tandem CAG-MerDreMer-Ki67p-RoxedCre plasmid.

### Generation of *αDKRC* transgene mice

Based on the results of the tandem CAG-MerDreMer-Ki67p-RoxedCre in cultured cells, a 5,442 bp αMHC-promoter was used to make a cardiomyocyte-specific αMHC-MerDreMer-Ki67p-RoxedCre (αDKRC). A single, tandem transgene was chosen to facilitate subsequent breeding strategies and avoid the Rosa26 locus, commonly used to generate transgenic reporters. Transgenic mice were generated by pronuclear injections of *C57BL/6J* oocytes using linearized *αDKRC* (Supplemental Figure 6B and C). Three independent founders were identified by polymerase chain reaction (PCR)-based genotyping across four regions of the ∼13 kb transgene to confirm the integrity of the transgene (Supplemental Figure 6C). Founder #1 did not transmit the αDKRC transgene to offspring. Founders were crossed to *C57BL/6J*, and genotype positive offspring that harbored αDKRC were treated with Tamoxifen (1 mg/kg IP daily for five days) or peanut oil vehicle (control) to test for the inducible expression of MerDreMer. Founder #2 produced genotype offspring; however, recombinant Dre protein was not expressed (data not shown). Hearts from mice derived from Founder #3 had Tamoxifen-dependent expression of MerDreMer localized to CM nuclei (Supplemental Figure 6D); therefore, *αDKRC* transgenic mice derived from Founder #3 were used for all subsequent experiments.

### *αDKRC/+::RLTG/RLTG* transgenic mice label adult CMs that reentered the cell cycle

A transgenic mouse harboring a CAG promoter-driven dual-recombinase responsive indicator *<u>R</u>ox-<u>L</u>ox-td<u>T</u>omato-e<u>G</u>FP* (*RLTG*) with a LoxP-STOP-LoxP cassette knocked into the Rosa26 locus^34^ was used to establish *αDKRC/+::RLTG/RLTG* mice and test the ability to label cycling adult CMs (Figure 1A). Transient administration of Tamoxifen to *αDKRC/+::RLTG/RLTG* mice were predicted to cause CM-specific expression of Dre recombinase and the removal of Rox-STOP-Rox cassettes from RLTG, resulting in the expression of tdTomato in CMs, and RoxedCre to produce Ki67-dependent Cre expression (Figure 1B). The approach has the unique advantage of requiring only transient exposure to Tamoxifen to “prime” the system for long-lasting effects, after which Cre is expressed in cycling CMs, as defined by activation of the Ki67 promoter. CM endoreplication was defined as single cytosolic GFP+ cells, indicative of cell cycle reentry without replication, and proliferation as two neighboring GFP+ CMs separated by cell membrane stained by wheat germ agglutinin (WGA).

Compared to treatment with peanut oil, Tamoxifen (1 mg/kg IP x 5 days) induced expression of tdTomato in CMs in *αDKRC/+::RLTG/RLTG* (Figure 1 C-F) and infrequent expression of eGFP in CMs (one eGFP+ CM identified in 6 ten-micron sections) (Figure 1G). Additional experiments were performed to examine the incorporation of BrdU into CMs of *αDKRC/+::RLTG/RLTG*. Adult *αDKRC/+::RLTG/RLTG* mice underwent implantation of an osmotic mini-pump for continuous BrdU administration. After a 7-day BrdU run-in period, mice were treated with Tamoxifen (1 mg/kg IP daily for two five-day intervals with a two day rest period in between intervals of Tamoxifen) (Supplemental Figure 7). After Tamoxifen administration, hearts were removed for immunofluorescence (IF) to identify eGFP+/BrdU+ CMs. Eighteen 10- micron sections throughout the hearts were examined, and 25 eGFP+ CMs were identified, of which 50% were BrdU+, consistent with the expression of Cre in CMs that reentered the cell cycle. The results support that *αDKRC/+::RLTG/RLTG* mice drive Cre in adult CMs that reenter the cell cycle.

### The *αDKRC* transgene is inserted in the intragenic region of Chromosome 14 between *MYH6* and *MYH7*

Transgenes inserted into the genome can potentially disrupt canonical cell cycle genes, confounding the interpretation of proliferation and endoreplication.^49^ Therefore, the locus of transgene insertion of *αDKRC* mice was identified by nested, inverse PCR. The αDKRC transgene was located in an intragenic region of Chromosome 14 between *MYH6* and *MYH7*, in an intron of *Myosin Heavy Chain Associated RNA Transcript* (*MHRT*), a long non-coding RNA implicated as a cardioprotective agent in the heart (Figure 2A and B).^50^ The expression of MYH6, MYH7, and MHRT was similar between CMs isolated from *αDKRC/+* mice and littermate controls (Figure 2C), suggesting that the αDKRC transgene did not disrupt neighboring gene expression. Heterozygous *αDKRC* transgenic mice were used for all the experiments.

**Figure 2.**
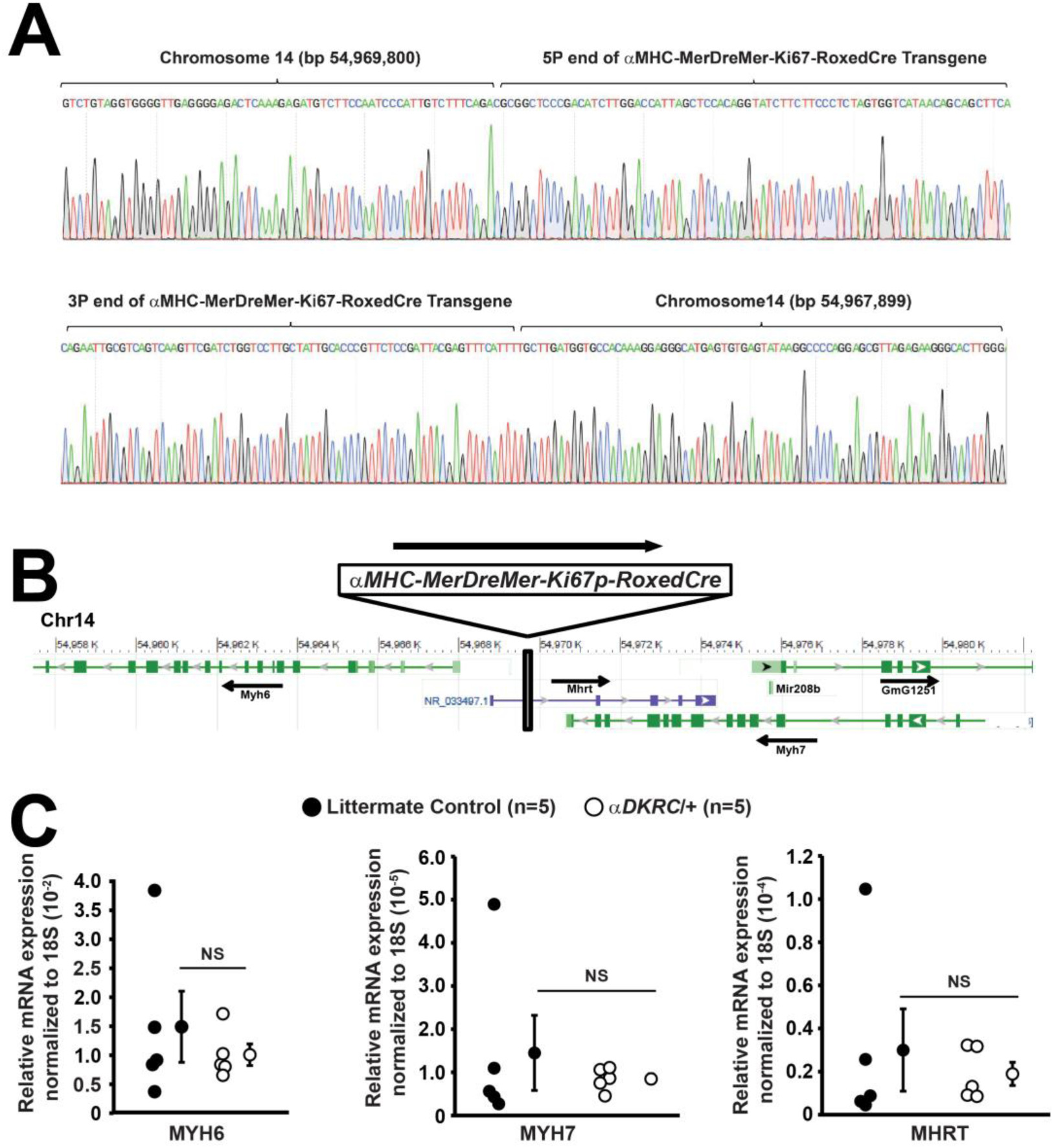
The αDKRC transgene is inserted in the intragenic regions of Chromosome 14 between *MYH6* and *MYH7*. (A) Representative DNA chromatograms after nested, inverse PCR of mice derived from *αDKRC* Founder #3. **(B)** The transgene is inserted between bp 54,967,899 and 54,969,800 of chromosome 14 based on Ensembl *Mus musculus* genome assembly version 97.28 (GRCm38.p6). Genomic map of the region of *αDKRC* transgene insertion. (**C)** Quantitate PCR (qPCR) of *MYH6*, *MYH7*, and *MHRT* expression from 12-week old hearts of *αDKRC* and littermate controls. N = 5 animals per group. Values a means ± SEM. mRNA expression normalized to 18S. No statistically significant differences (NS) between groups using Student’s t-test.

### *αDKRC/+::RLTG/RLTG* label cycling CMs after Ischemia-Reperfusion MI

Few CMs reenter the cell cycle in response to injury; however, detection of these rare events is a challenge for the field as a whole because cell cycle reporters alone are inadequate to define proliferating CMs because increased ploidy (i.e., endoreplication) can occur without proliferation.^28^ However, the *αDKRC/+::RLTG/RLTG* mice were designed to provide a readout of CM cycling events, integrated over time, starting at brief Tamoxifen exposure, and spatially across the myocardium. CM endoreplication was defined as single GFP^+^ cells, indicative of cell cycle reentry without replication, and proliferation as two neighboring GFP^+^ CMs separated by cell membranes stained by WGA.

Sham and I/R MI surgeries were performed in *αDKRC/+::RLTG/RLTG* mice and subsequent, CM cell cycle events were quantified in the infarct, border, and remote zones of the injured myocardium. *αDKRC/+::RLTG/RLTG* mice were treated with Tamoxifen, to activate the reporter system, followed by a two week recovery period before undergoing 60 minutes of LAD ligation ischemia followed by reperfusion or sham surgeries. Two weeks after I/R MI, single endoreplicated and paired proliferated eGFP+ CMs were readily identified (Figure 3A-D). GFP^+^ CMs increased in number after I/R MI compared to sham (5.8 ± 0.5 vs. 3.3 ± 0.3 CMs per ten-micron section, I/R MI vs. Sham, p<0.01) with the majority of single eGFP+ CMs (74.0 ± 3.9%) and paired eGFP^+^ CMs (89.3 ± 7.3%) localized in the border zones after I/R MI (Figure 3E). The ratio of endoreplication (ploidy) to proliferation was ∼9 to 1 (5.2 ± 0.4 single vs. 0.6 ± 0.2 paired CMs) in the I/R MI group (Figure 3F). Moreover, paired eGFP+ CMs were more abundant in the border zones compared to infarct and remote zones (12.0 ± 2.5%, 7.0 ± 4.4%, and 5.8 ± 5.0% for the border, infarct, and remote zones, respectively), suggesting that cycling CMs are more likely to undergo polyploidy than replication. In additional experiments, we implanted EdU minipumps and observed similar labeling of GFP+ CMs using Click-It chemistry (Supplemental Figure 8).

**Figure 3.**
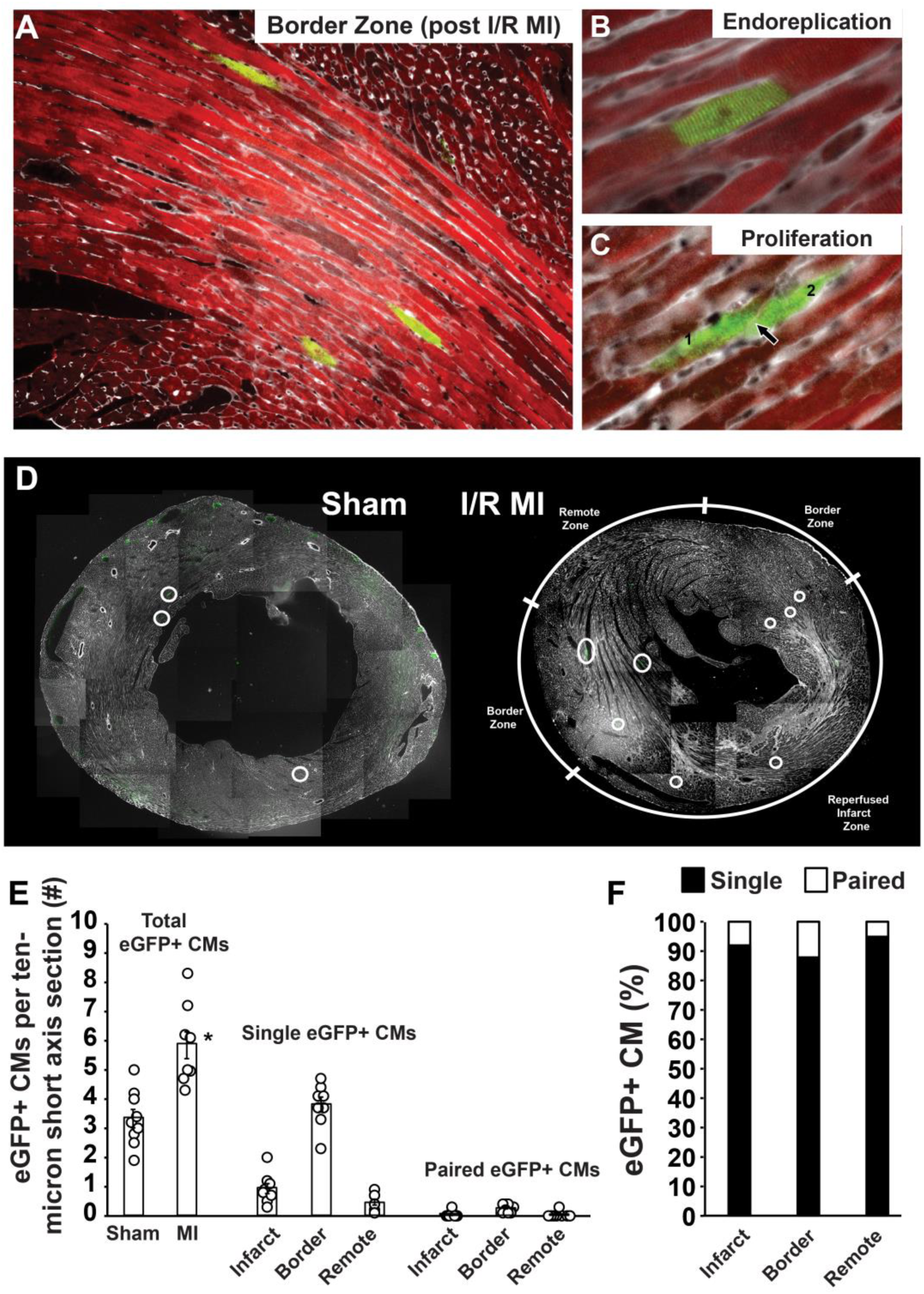
Cycling CMs increase after I/R MI and predominantly endoreplicate. Discrimination between CM endoreplication and proliferation using *αDKRC::RLTG* mice. (A) Representative image showing individual eGFP^+^ CMs in the border zone of the infarct. Red = anti-tdTomato antibody, Green – anti-eGFP antibody, and Gray – WGA. (**B and C)** Representative examples of a single eGFP^+^ CM **(B)** and two neighboring eGFP^+^ CMs with cell boundary between the cells labeled by WGA **(C)**. The arrow denotes WGA stating between two eGFP^+^ CMs **(D)** Representative short-axis IFH images of hearts from *αDKRC::RLTG* mice two weeks after Sham (left) or I/R MI (right). White circles denote eGFP+ CMs. **(E)** Quantification of eGFP^+^ CMs from sham and I/R MI hearts, and the infarct, border, and remote zones of I/R MI hearts. *p<0.05 for MI vs. Sham by t-test. N = 6-9 animals per group, n = 16-24 sections per animal. Open circles are values for individual animals, bar are group means ± SEM. **(F)** Ratio of single to paired eGFP+ CMs after I/R MI corresponding to panel B.

### The ablation of endogenously cycling adult CMs worsens left ventricular function after I/R MI

To date, the vast majority of investigations of adult CM cycling has examined the effects of ablating or overexpressing specific genes in CMs using *α*MHC-Cre, *α*MHC-MerCreMer, cTNT-Cre, or other “pan”-CM drivers. Certainly, the enhanced cycling globally in all CMs has been associated with improved myocardial function after injury. However, to the best or our knowledge, no investigation has ablated endogenously cycling adult CMs and asked “does loss of cycling CMs impact myocardial function after injury?” *αDKRC* provides the unique opportunity to investigate this important question. Since Cre recombinase expression is restricted to adult cycling CMs in *αDKRC* mice, we made *αDKRC::DTA* mice to ablate cycling CMs and interrogate the effects of loss of these cells on cardiac function after I/R MI (Figure 4 A and B). Male and female *αDKRC::DTA* and littermate controls (*+::DTA*) underwent pre-treatment with Tamoxifen or Peanut oil before a recovery period, followed 60 minute I/R MIs (Figure 4C). *αDKRC::DTA* pre-treated with Tamoxifen to ablate cycling CMs had increased end-systolic volumes and worsened left ventricular ejection fractions four weeks after I/R MI compared to peanut oil-treated and *+::DTA* controls (Figure 4D). For any given infarct size, the LVEF was worse in *αDKRC::DTA* pre-treated with Tamoxifen compared to all other control groups (Figure 4E-G). The data suggest that ablation of adult cycling CMs worsens myocardial function after injury.

**Figure 4.**
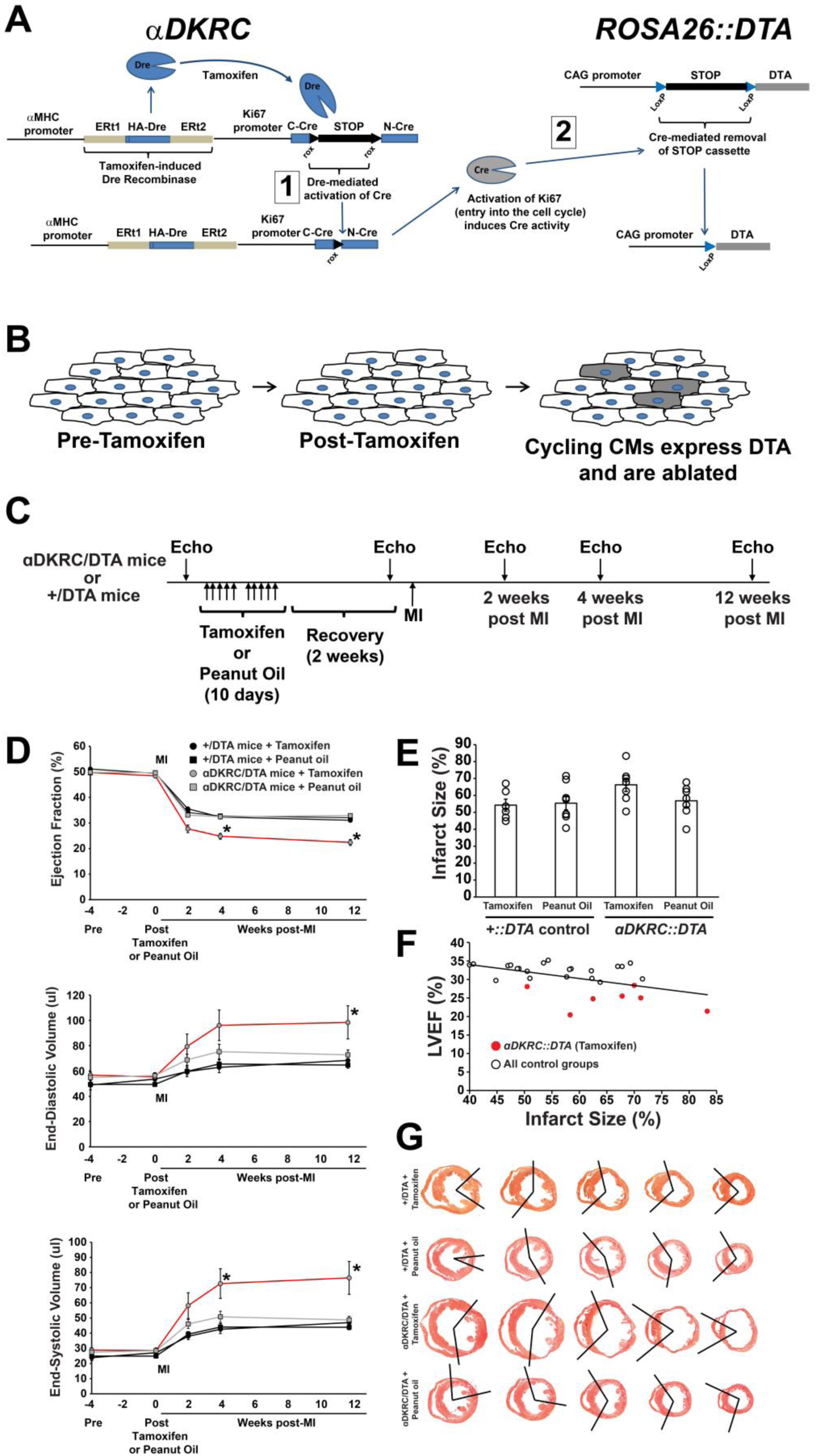
Endogenously cycling adult cardiomyocytes contribute to myocardial function after injury. (A and B) *αDKRC::Rosa26-Diphtheria Toxin (DTA)* (*αDKRC::DTA)* mice were generated to selectively ablate adult cardiomyocytes the reentered the cell cycle. Pre-treatment with Tamoxifen “primes” Cre recombinase to be expressed in CMs that have activated Ki67. Then, Cre recombinase excises a STOP cassette, causing the expression of DTA and ablation of cycling CMs. **(C)** Protocol using *αDKRC::DTA* and +*::DTA* littermate controls treated with Tamoxifen (1 mg/kg IP daily x five days x two cycles) or Peanut Oil (control) followed by recovery period before undergoing 60 minutes of LAD-ligation ischemia and reperfusion (I/R MI). Serial echocardiography was performed prior to and after MI. **(D)** Left ventricular Ejection Fraction (%), Left Ventricular End-systolic and End-diastolic volumes (ul) showing worsened LV chamber sizes and function in *αDKRC::DTA* mice pre-treated with Tamoxifen compared to controls. N= 7-9 animals per group. Values are means ± SEM. *p<0.01 by ANOVA followed by Bonferroni post hoc test correction for multiple comparisons. **(E)** Infarct sizes for each group corresponding to Panel D. N= 7-9 animals per group. Open circles are values for individual animals, bar are group means ± SEM. **(F)** Relationship between Left ventricular Ejection Fraction (%) and infarct sizes (bottom panel). White dots are all control groups. Red dots are *αDKRC::DTA* mice pre-treated with Tamoxifen. **(G)** Representative short axis sections of groups corresponding top Panel D from base (left) to apex (right) stained with PicroSirius Red. Infarcted regions are denoted by black bars.

## Discussion

Endoreplication occurs when a cell proceeds through the four canonical phases of the cell cycle.^51^ However, cytokinesis is incomplete, producing bi-nucleated or mono-nucleated polyploid cells. Polyploidy occurs among species during critical windows of healthy growth and in response to disease.^28, 51–53^ Moreover, polyploidy has been proposed to be one mechanism limiting the regenerative capacity of mammalian organs to respond to injury.^53, 54^ Mammalian CMs undergo endoreplication and binucleation during postnatal heart growth, in response to hypertension, and as a result of injury.^17, 28, 52^

The accurate detection and quantification of endoreplication and proliferation *in vivo* are critical to interrogating signaling pathways that potentially influence the fate of cycling cells. This is particularly important when assessing adult CM responses to injury. Cell cycle marker expression, usually in conjunction with nucleotide incorporation, has been used to quantify CM cycling events. The expression of Ki67, Aurora B, and Histone H3 phosphorylation mark cells that enter the cycle but cannot distinguish between endoreplicating and proliferation. However, estimates of cycling CMs based on marker expression suggest ranges of 0 to 0.8% and 0.01 to 3.8% for uninjured and injured myocardium, respectively.^7, 13, 55^ Measurements of nucleotide analog incorporation using BrdU, Edu, or ^13^N-thymidine into mammalian CMs are reported in the range of 0 to 0.6% in uninjured models, and 0.015 to 3.2% after injury.^3, 7, 21, 55–59^ Using aDKRC and BrdU, we estimated ∼0.07% and ∼0.02% eGFP+ CMs per total CMs per section for I/R MI and sham groups, respectively; comparable to published estimates of cycling CMs based on incorporation of BrdU or EdU and expression of markers of the cell cycle. ^3, 7, 21, 55–59^

The dose, route of administration, and duration of exposure vary among published investigations. The disruption of vascularity that occurs in the setting of injury potentially limits the access of BrdU to regions of the myocardium, confounding interpretation of cycling cells. Moreover, the measurement of nucleotide analog incorporation occurs during S-phase and cannot distinguish between endoreplication (polyploidy) and proliferation of CMs. Even combining these methods, colocalization of markers to CMs can be challenging.

Although analyses of marker expression and thymidine analog incorporation are valuable methods to detect cycling cells, particularly in large animals and human pathology, innovative approaches that use transgenic mice lead to greater mechanistic insights into CM cycling and proliferation, particularly after injury. Transgenic mice such as *Ki67-RFP* and *Ki67-^iresCreER^* mice label cycling cells using RFP or tamoxifen-inducible Cre.^36, 37^ However, Ki67-RFP labels all actively cycling cells, lacking cell type-specificity and the ability to integrate cycling.^37^ *Ki67- ^iresCreER^* drives a tamoxifen-inducible Cre in dividing cells and could be used with a Cre-inducible, CM-specific fluorescent reporter to label cycling CMs.^36^ A *Ki67-^iresCreER^* strategy would require continuous exposure to Tamoxifen to activate the system, potentially confounding interpretation because of the known effects of prolonged tamoxifen on cardiac function.^60, 61^ A recent eGFP-Anillin reporter mouse distinguishes between cell cycle events, at the time of cell cycling, based on localization of the reporter to the midbody during cellular division.^62^ Fluorescent-based cell cycle indicators (Fucci) label actively cycling cells, particularly during the postnatal transition when CMs exit the cell cycle (Supplemental Figure 9).^41, 63^ Additionally, Fucci-based transgenics combined with thymidine analogues can identify cells that re-enter the cell cycle after injury (Saucerman and Wolf, *unpublished results*); however, this combination alone still cannot distinguish between endoreplication and proliferation. By comparison, our *αDKRC/+::RLTG/RLTG* mice label adult CMs, require a limited Tamoxifen treatment to achieve durable reporter expression, and integrate cycling events to facilitate improved detection of low-occurrence cycling events over prolonged periods of time.

Similar to our *αDKRC* mice, transgenic mosaic analysis with double markers (MADM) mice label clones of dividing cells based on Cre-mediated recombination during G2 phase^20, 64^. The labeling efficiency of MADM mice is low because of Cre-loxP-dependent inter-chromosomal mitotic recombination, requiring recombination to occur on two separate chromosomes, to generate labeled daughter cells. The use of MADM to detect adult cycling CMs requires an inducible Cre (i.e., *αMHC-MerCreMer*)^20^ and the continuous presence of Tamoxifen to activate the system, a potential confounder because Tamoxifen is known to cause a cardiomyopathy.^60, 61^ Moreover, MADM cannot drive Cre specifically in cycling CMs. Additional approaches include CM-specific expression of BrainBow or Confetti reporters where Cre-mediated recombination of multiple fluorescent reporters labels the clonal expansion of CMs.^15^ Similar to Confetti-based strategies, CM endoreplicative and proliferative events can be inferred based on the labeling of single and paired eGFP+ CMs using *αDKRC/+::RLTG/RLTG* mice. However, an important distinction of *αDKRC/+* is the postnatal expression of inducible Cre that is restricted to actively cycling adult CMs, thereby providing a means to interrogate the contribution of cycling CMs after injury to myocardial function and homeostasis. Our results using *αDKRC/+::RLTG/RLTG* mice and examining cycling events across multiple short-axis sections of the myocardium suggest that endoreplication was the predominant outcome of cycling CMs, with a ratio of 9:1, single to paired eGFP positive CMs.

There are potential unlikely and rare limitations of *αDKRC/+::RLTG/RLTG* mice. First, the activation of the 1.5 bp Ki67 promoter defines “cycling”. Although unlikely, a CM that enters the cycle may fails to proceed through S-phase, leading to an overestimation of cycling events, as observed in cells that overexpress p53 or p21.^65^ Second, while rare, tamoxifen may not uniformly activate the αDKRC reporter across the entire myocardium, resulting in an underestimation of cycling CMs. Third, a pair of eGFP positive CMs in the hearts of *αDKRC/+::RLTG/RLTG* mice could represent neighboring endoreplicating CMs, thereby over-estimating CM proliferation. However, the expression of eGFP in two neighboring CMs by chance alone is predicted to be an infrequent event. For example, the random occurrence of two neighboring CMs in a square field of 1000 cells, assuming four edges per cell is ∼0.35%, the estimate decreasing as the field of cells increases (Supplemental Figure 10). Fourth, the measurements of eGFP+ CMs were made from multiple 10 micron thin sections (Supplemental Figure 3). Therefore, neighboring eGFP+ CMs above or below the plane of section could be potentially missed, underestimating the quantification of proliferating CMs. Fifth, the daughter cells of proliferating CMs could die or migrate away from the cycling CM, underestimating the number of CM replicating events. Sixth, if Dre excised both LoxP and Rox-flanked cassettes, then a false-positive eGFP^+^ CM would be produced, potentially overestimating CM cycling. However, dual Dre and Cre approaches are rigorously specific enough for lineage-tracing transgenic models ^31, 32, 34, 35^ and our validation experiments in cell culture supported the recombinase specificities required to create the αDKRC mouse. Moreover, we observed an increase in eGFP+ CMs after injury, particularly in the border zones, consistent with CM cycling. Finally, the RLTG and DTA transgenes are driven by CAG promoters comprised of a cytomegalovirus early enhancer element promoter, the first exon and the first intron of chicken beta-actin gene, and the splice acceptor of the rabbit beta-globin gene.^66^ The CAG promoter can be downregulated in some cell types.^67–69^ In the context of RTLG and DTA, a silencing of the CAG promoter could potentially lead to an under-estimation of cycling or inefficiency of ablation, respectively. However, we observed robust tdTomato expression (Figure 1 E-G), consistent with functional CAG promoter expression in CMs. Moreover, the issues with CAG promoters also apply to many reporters, including MADM and Confetti/BrainBow.

Although considered a “gold-standard”, thymidine analog incorporation into cycling cells *in vivo* has limitations. The 50% of BrdU-, eGFP+ CMs observed *αDKRC/+::RLTG/RLTG* mice may represent inefficient BrdU uptake, incomplete unmasking of BrdU epitopes, difficulties identifying co-localization, or problems of antibody access to the labeled DNA, factors known to confound BrdU labeling experiments. Lastly, the αDKRC technology cannot determine whether proliferating CMs cells originated from mono- or bi-nucleated CMs, but this does not prevent interpretation of proliferation and endoreplication.

Increased cycling of CMs was observed after I/R MI, particularly in the border zone, compared to sham control, consistent with published literature.^13^ The majority of published literature uses permanent LAD ligation MI models.^4, 13, 22, 24, 55^ The I/R MI model more closely approximates the clinical scenario of MI with successful reperfusion and produces “reperfused infarct” (i.e., area-at-risk + infarct), border, and remotes zones (Supplemental Figure 3B).^70^ The “reperfused infarct” zone contains non-transmural scar and surviving myocytes. I/R MI, at a precisely controlled body temperature of 37.5°C based on our experience, allows a more detailed regional characterization of cycling CMs than the permanent ligation MI model. Compared to permanent ligation MI, the reperfused infarct has injured CMs that experienced hypoxia, a stimulus shown to stimulate potential cycling.^6, 7^ The reperfused infarct has increased vascularity and less fibrosis based on the non-transmural nature of the infarct.

*αDKRC/+* builds on the established transgenics that employ *α*MHC promoters (Supplemental Figure 11). Importantly, *αDKRC /+* mice not only label adult cycling CMs but also provide opportunities to ablate or express genes specifically in these subpopulations. Using *αDKRC::DTA* mice, cycling CMs were ablated and cardiac function worsened after I/R MI, suggesting that these cells contribute to myocardial function. Prior investigations using 5-fluorouracil to blunt general cell proliferation, including CMs, showed that cardiac cell proliferation was required for protection against ischemia-reperfusion injury, but not necessary for exercise-induced cardiac growth.^71^ A major limitation of 5-fluorouracil is the non-specific effects, thereby potentially blunting angiogenesis, fibroblast/myofibroblast proliferation, and inflammatory cells. *αDKRC::DTA* mice specifically target cycling CMs, thereby avoiding the non-specific effects of chemical inhibitors. The observations that cardiac function progressively worsened after I/R MI in *αDKRC::DTA* mice was unexpected because cycling CMs are scare and assumed to not significantly impact cardiac function. One potential mechanism to explain the observations is that cycling CMs express paracrine factors involved in the injury response, and loss of these factors worsens myocardial function. This requires thinking of subpopulations of cycling CMs as secretory, and not exclusively contractile, cells. If this is the case, then a few cells can have significant impact over a larger area of the myocardium and a modest increase in cycling cells can have a greater influence on myocardial function than that expected if simply ascribing contractile function to this subpopulation of CMs. Future investigations will use the *αDKRC* mice to molecularly characterize how cycling CMs differ from the vast majority of non-cycling adult CMs and if paracrine factors control myocardial health after injury, exercise, and aging.

### Study Approval

All animal care and surgeries were in accordance with UVA ACUC Policy on Rodent Surgery and Perioperative Care under ACUC-approved animal protocol (UVA ACUC Wolf Lab protocol #4080).

## Author Contributions

LAB, AY, HOB, and MJW designed experiments, generated reagents, performed and analyzed experiments, and wrote the manuscript. AY and HOB performed rodent surgeries. MJW designed the transgenic mice and secured funding for the project.

## Acknowledgements

**Sources of Funding:** This work was solely supported by the University Of Virginia Center Of Excellence in Cardiovascular Genetics.

## Supplemental Figures

**Supplemental Figure 1.**
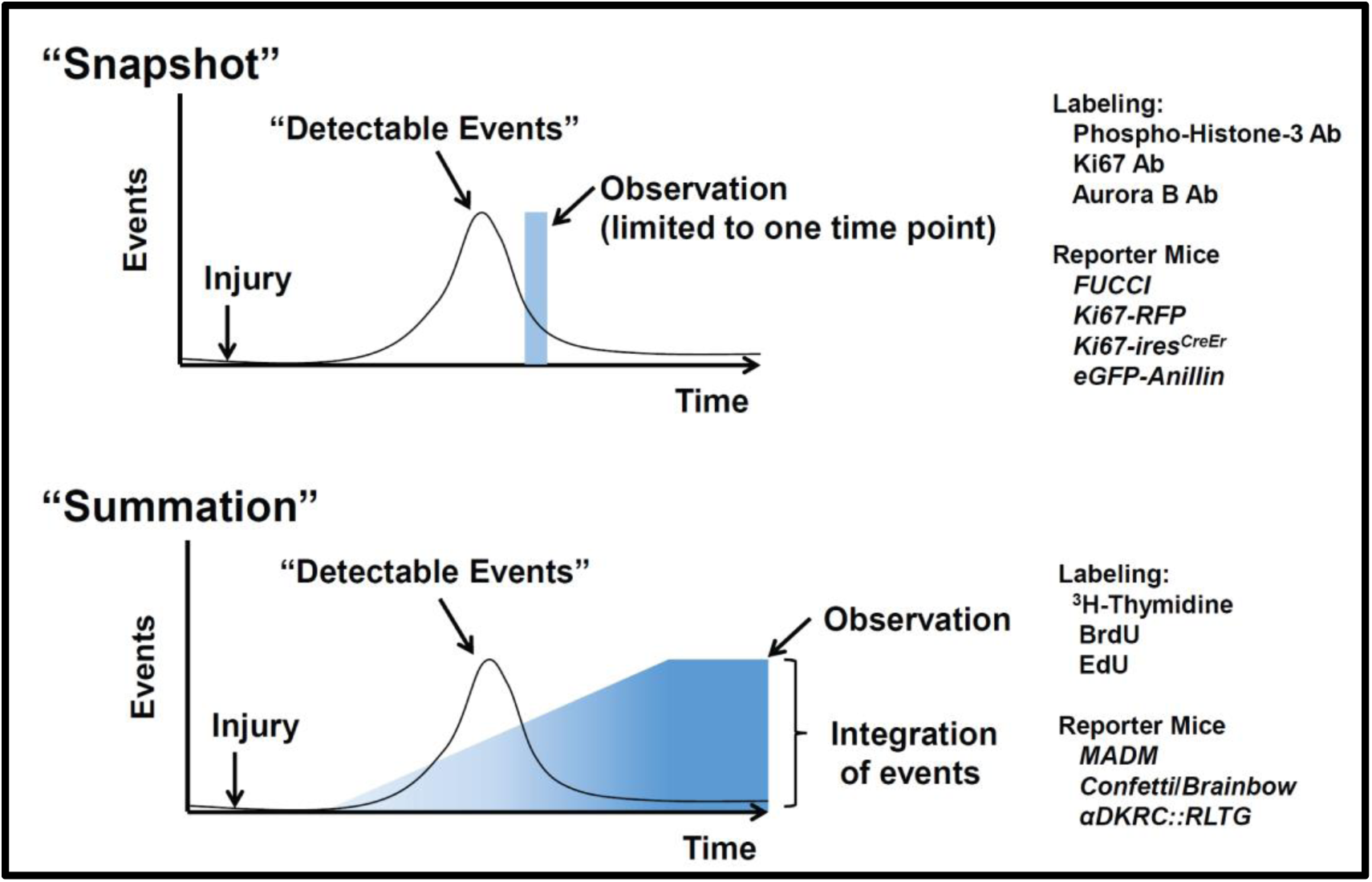
The “Snapshot” and “Summation” approaches to observing cycling events. “Detectable events” such as cycling cells occur after an injury or stimulus. The “Snapshot” view (Top) relies on observations, such as the expression of phospho-Histone-3, Aurora B, Ki67, or fluorescence-based reports like FUCCI, at a single time point. The ‘Summation” view (Bottom) relies on the incorporation and retention of a marker, such as BrdU, ^3^H-Thymidine, or EdU, reporting the integrative signal of “detectable events” over periods of time.

**Supplemental Figure 2.**
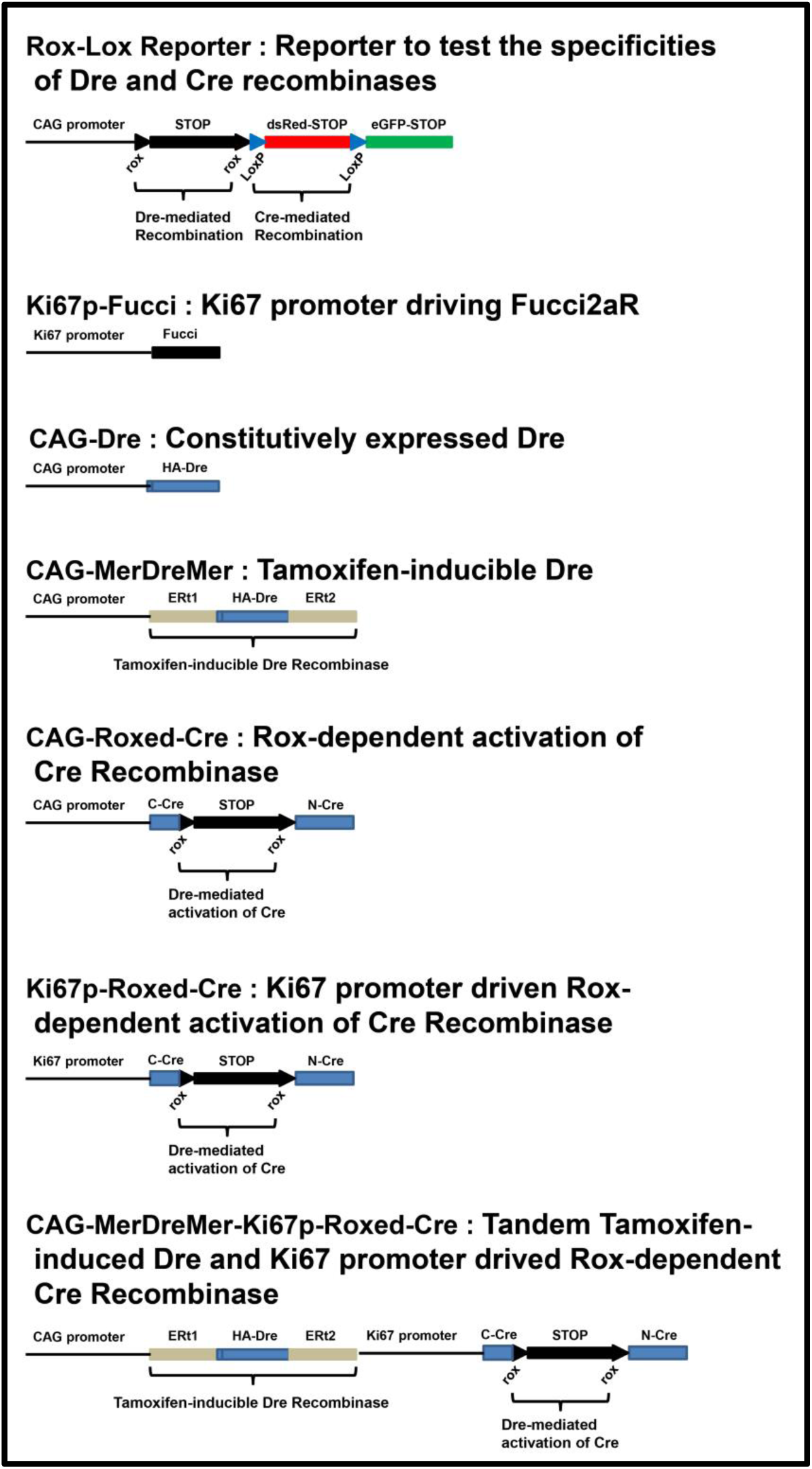
DNA constructs used to validate Tamoxifen-inducible Dre, RoxedCre, and mouse Ki67 promoter.

**Supplemental Figure 3:**
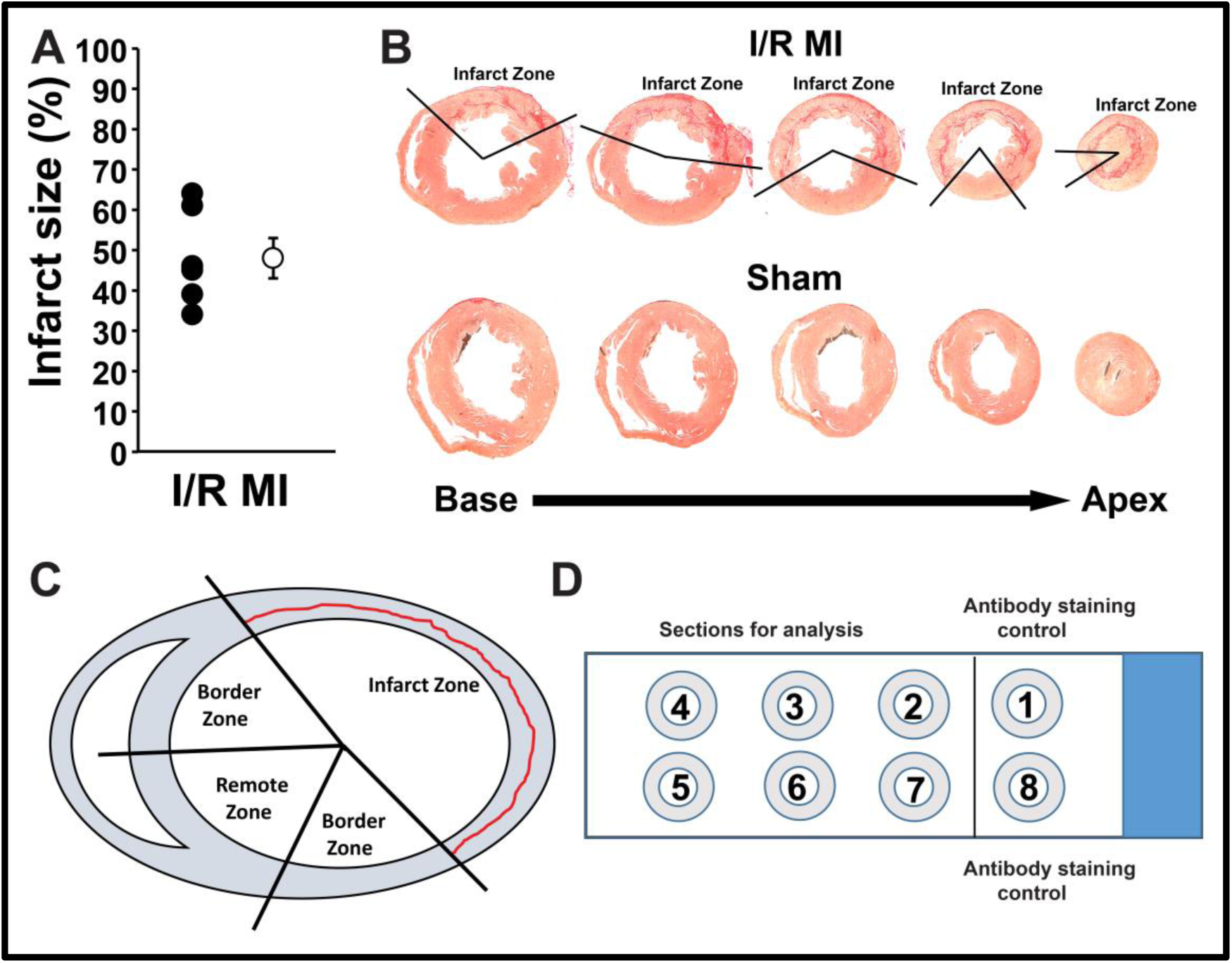
Definition of I/R MI zones and sections. A, Summary of infarct sizes of *αDKRC::RLGT* mice two weeks after I/R MI. N = 6 animals. Values for individual mice (black circles) and the mean ± SEM (white circle) are shown. Infarct sizes were calculated as the percentages of myocardium averaged across the middle three short-axis sections per mouse. **B,** Representative short axis sections of hearts of I/R MI and Sham animals stained with Picrosirus Red. Infarct zones are denoted. **C,** Schematic of the infarct, remote, and border zones. **D,** Schematic of a slide containing 8 ten-micron short-axis sections of a heart used for immunofluorescence. Sections #1 and #8 are antibody controls. Sections #2 through #7 are used to quantify eGFP^+^ CMs.

**Supplemental Figure 4.**
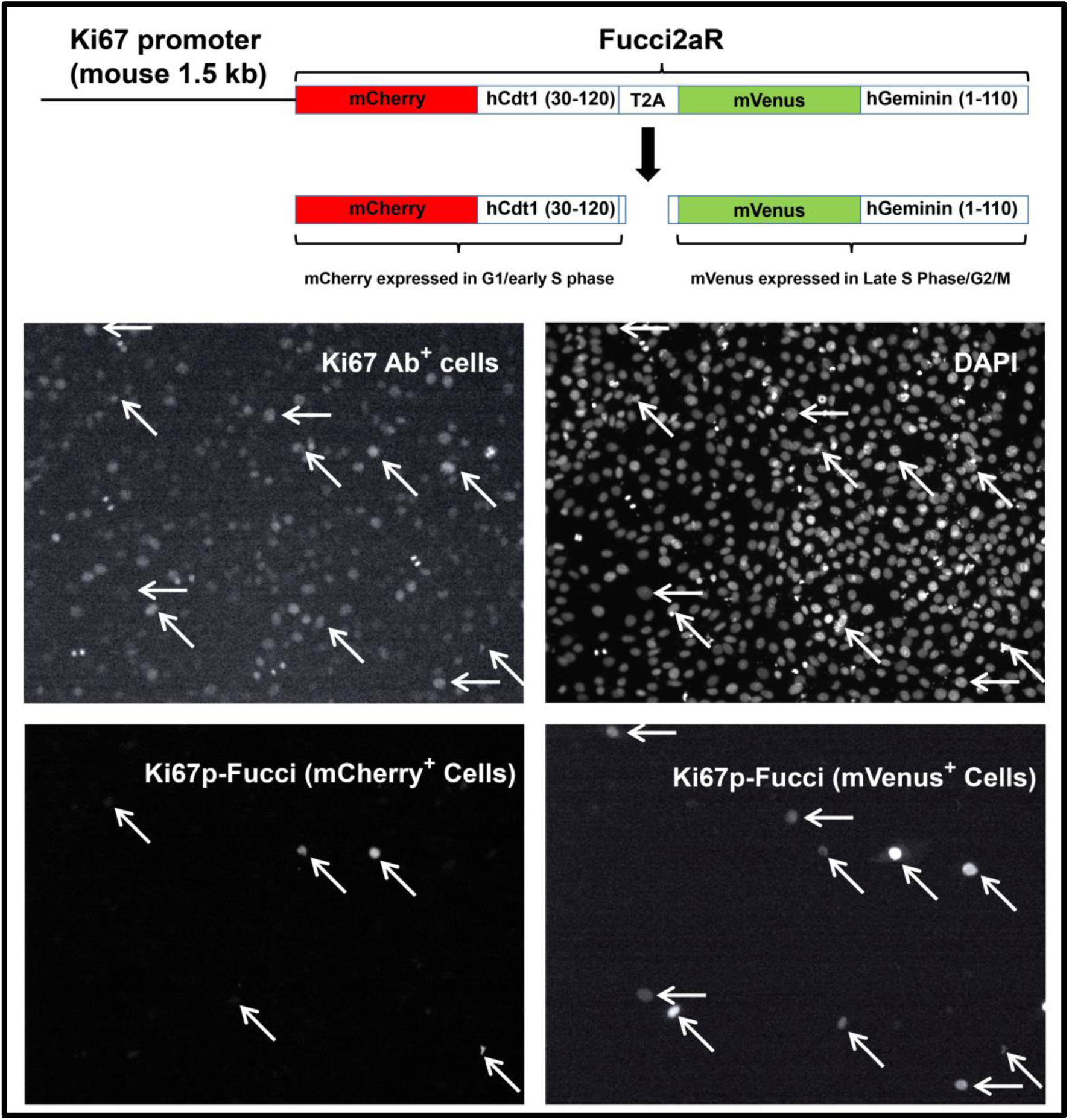
Validation of mouse Ki67 promoter. Top, The 1.5 kb Ki67 promoter described by Zambon was isolated from *C57B6J* genomic DNA, validated by DNA sequencing, and used to drive a fluorescent ubiquitin-based cell cycle indicator (Fucci2aR). Bottom, HEK293T cells were transfected with 1 ug of Ki67p-Fucci2aR plasmid using Lipofectamine. After 48 hours, the cells were fixed in 4% PFA, permeablized using Triton-X, and examined using immunofluorescence after staining with anti-Ki67, anti-mCherry, and anti-GFP antibodies and DAPI. White arrows label cells that expressed Fucci2aR and co-localized with an anti-Ki67 antibody. Ki67 expression co-localized with Fucci positive cells.

**Supplemental Figure 5.**
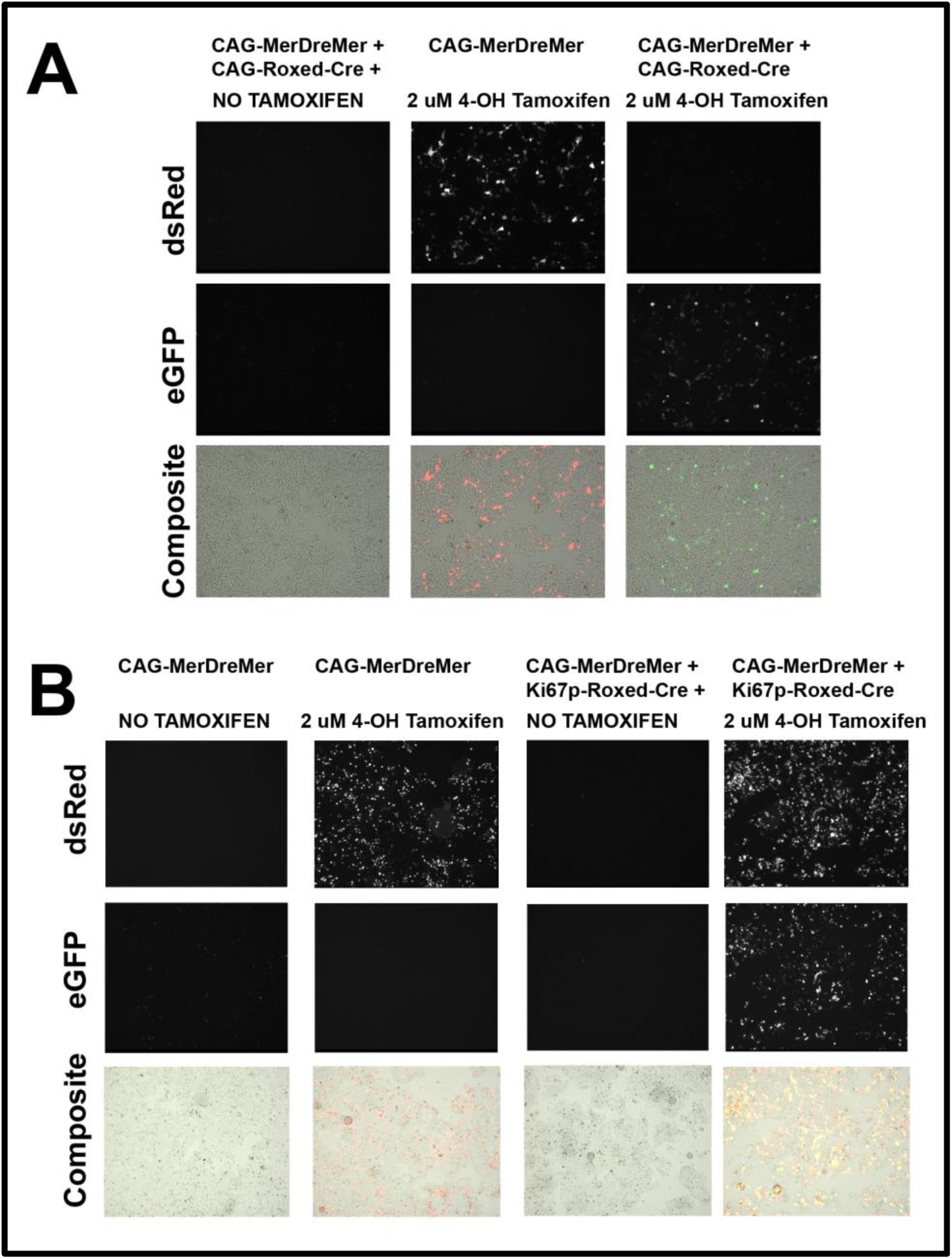
Validation of Tamoxifen-inducible Dre, RoxedCre, and mouse Ki67 promoter in HEK-293T cells. HEK293T cells were transfected with CAG-Rox-STOP-Rox-LoxP-dsRed-STOP-LoxP-eGFP-STOP reporter and combinations of CAG-MerDreMer, and CAG-RoxedCre **(A)** or Ki67p-RoxedCre **(B)**. After 48 hours, cells were treated with vehicle or 2uM 4-OH Tamoxifen for 24 hours, and the expression of live dsRed and eGFP was visualized to confirm the specificity of Tamoxifen-inducible Dre and RoxedCre.

**Supplemental Figure 6.**
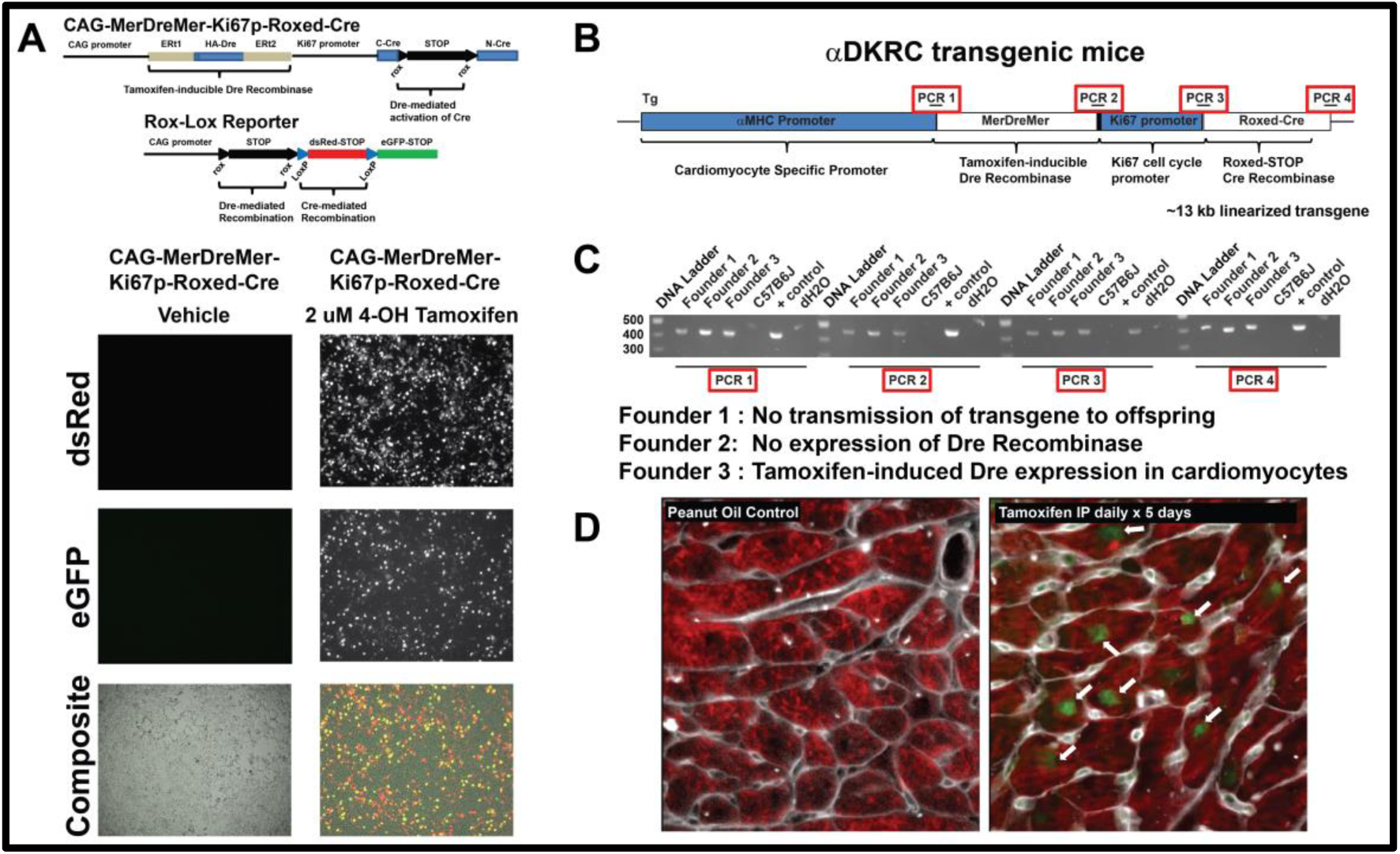
Validation of Tamoxifen-inducible Dre recombinase and cell cycle-dependent Ki67p-RoxedCre recombinase, and creation of *αDKRC* transgenic mice. A, Top, CAG-MerDreMer-Ki67p-RoxedCre and CAG-Rox-STOP-Rox-LoxP-dsRed-STOP-lox-eGFP-STOP reporter constructs. Bottom, Representative live images of HEK293T cells transfected with tandem CAG-MerDreMer-Ki67p-RoxedCre and CAG-Rox-STOP-Rox-LoxP-dsRed-STOP-lox-eGFP-STOP reporter and treated with 2uM 4-OH Tamoxifen for 48 hours. **B,** Linearized αDKRC transgene structure. Red boxes denoting PCR1, PCR2, PCR3, and PCR4 correspond to the regions of the transgene used for subsequent genotyping. **C,** Agarose gel of PCR genotyping of 3 independent *αDKRC* founder mice. Founder #3 was used for all subsequent experiments. **D,** Representative immunofluorescence images of *αDKRC* transgene mice derived from Founder #3 treated with vehicle (Peanut Oil) or Tamoxifen (1 mg/kg IP daily x 5 d). Paraffin sections were stained for cardiac troponin (Red), HA-Dre (Green), and Wheat Germ Agglutinin (WGA) (Gray). White arrows point to Dre localized to CM nuclei.

**Supplemental Figure 7.**
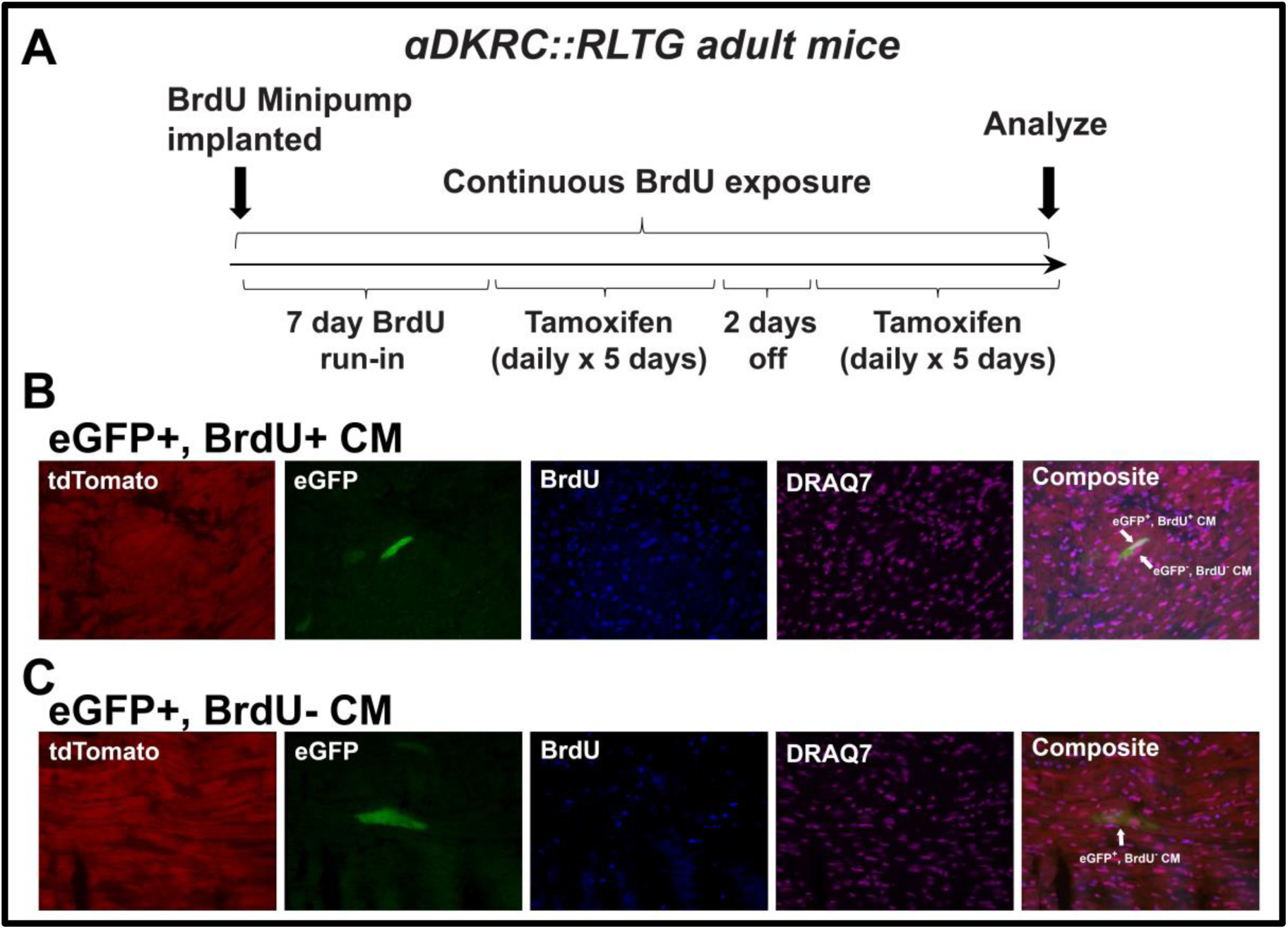
Co-localization of BrdU with eGFP CMs in *αDKRC::RLTG*. A, Twelve-week old *αDKRC::RLTG* underwent implantation of an osmotic minipump contacting BrdU. After a seven day run-in period, Tamoxifen was delivered (1 mg/kg IP daily for five days x 2 cycles with a 48 hour rest period), and hearts were isolated for paraffin embedding and immunofluorescence. Twenty-five eGFP+ CMs with DRAQ7+ nuclei were examined for the presence of BrdU from eighteen 10-micron sections of whole hearts in the short axis per animal were examined (N=3 animals). Twelve of the twenty-five eGFP+ CMs were BrdU+ (50%). **B and C,** Representative immunofluorescence of sections stained with anti-tdTomato, ant-eGFP, anti-BrdU, and DRAQ7. White arrows show eGFP+, BrdU+ and eGFP+, BrdU-CMs.

**Supplemental Figure 8.**
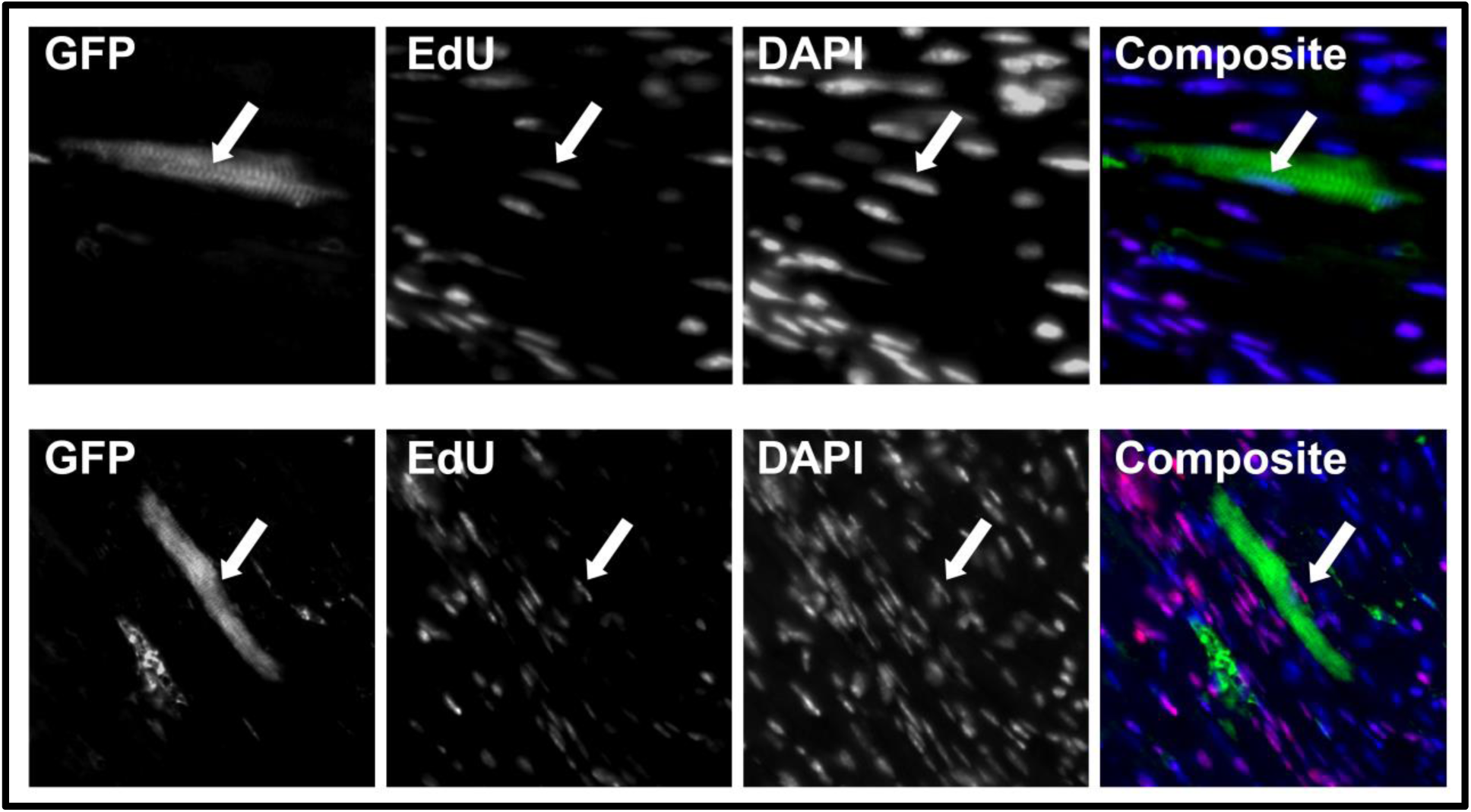
Co-localization of EdU with eGFP CMs in *αDKRC::RLTG*. A, Twelve-week old *αDKRC::RLTG* underwent two cycles of Tamoxifen administration (1 mg/kg IP daily x 5 days) over two weeks followed by a two-week recovery before undergoing 60 minutes of ischemia by LAD ligation 1 mm below the left auricle followed by reperfusion. An osmotic minipump contacting EdU was implanted immediately following MI surgery. Two weeks after MI, hearts were isolated for paraffin embedding and immunofluorescence and Click-It chemistry to detect EdU. Representative GFP+/EdU+ CMs in the border zones are shown.

**Supplemental Figure 9.**
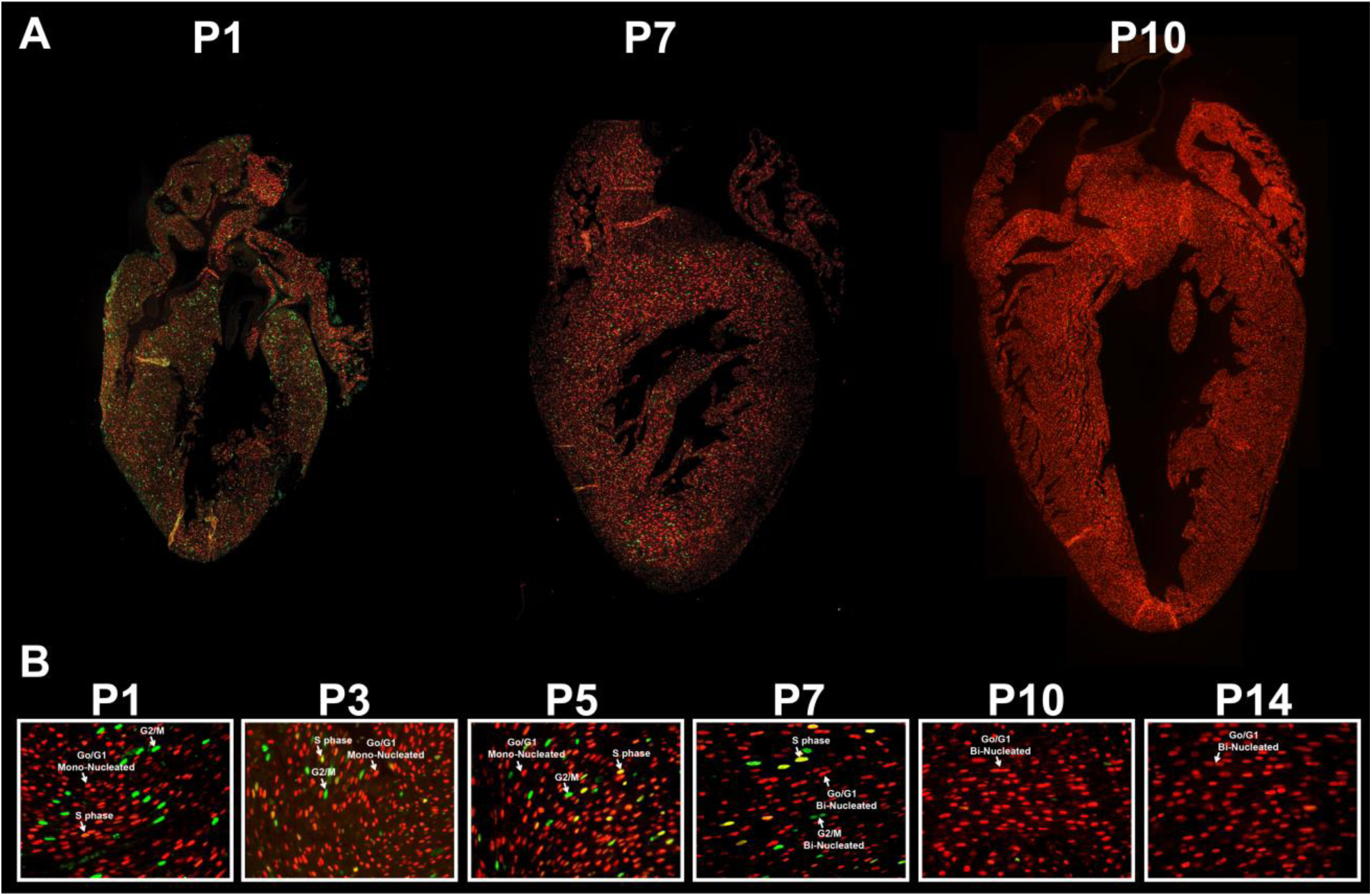
Cycling CMs of *αMHC-Cre/+::CAG-STOP-Fucci2aR/CAG-STOP-Fucci2aR*. (A) Composite long-axis sections of hearts from P1, P7, and P10 *αMHC-Cre/+::CAG-STOP-Fucci2aR/CAG-STOP-Fucci2aR* mice showing G_0_/G_1_ (mCherry**^+^**/mVenus**^-^**), S (mCherry**^+^**/mVenus**^-^**), and G_2_/M (mCherry**^-^**/mVenus**^+^**) CMs. **(B)** 40x magnification of myocardium from P1, P3, P5, P7, P10, and P14 *αMHC-Cre/+::CAG-STOP-Fucci2aR/CAG-STOP-Fucci2aR* mice showing G_0_/G_1_ (mCherry**^+^**/mVenus**^-^**), S (mCherry**^+^**/mVenus**^-^**), and G_2_/M (mCherry**^-^**/mVenus**^+^**) during the postnatal period when CMs undergo binucleation and exit the cell cycle.

**Supplemental Figure 10.**
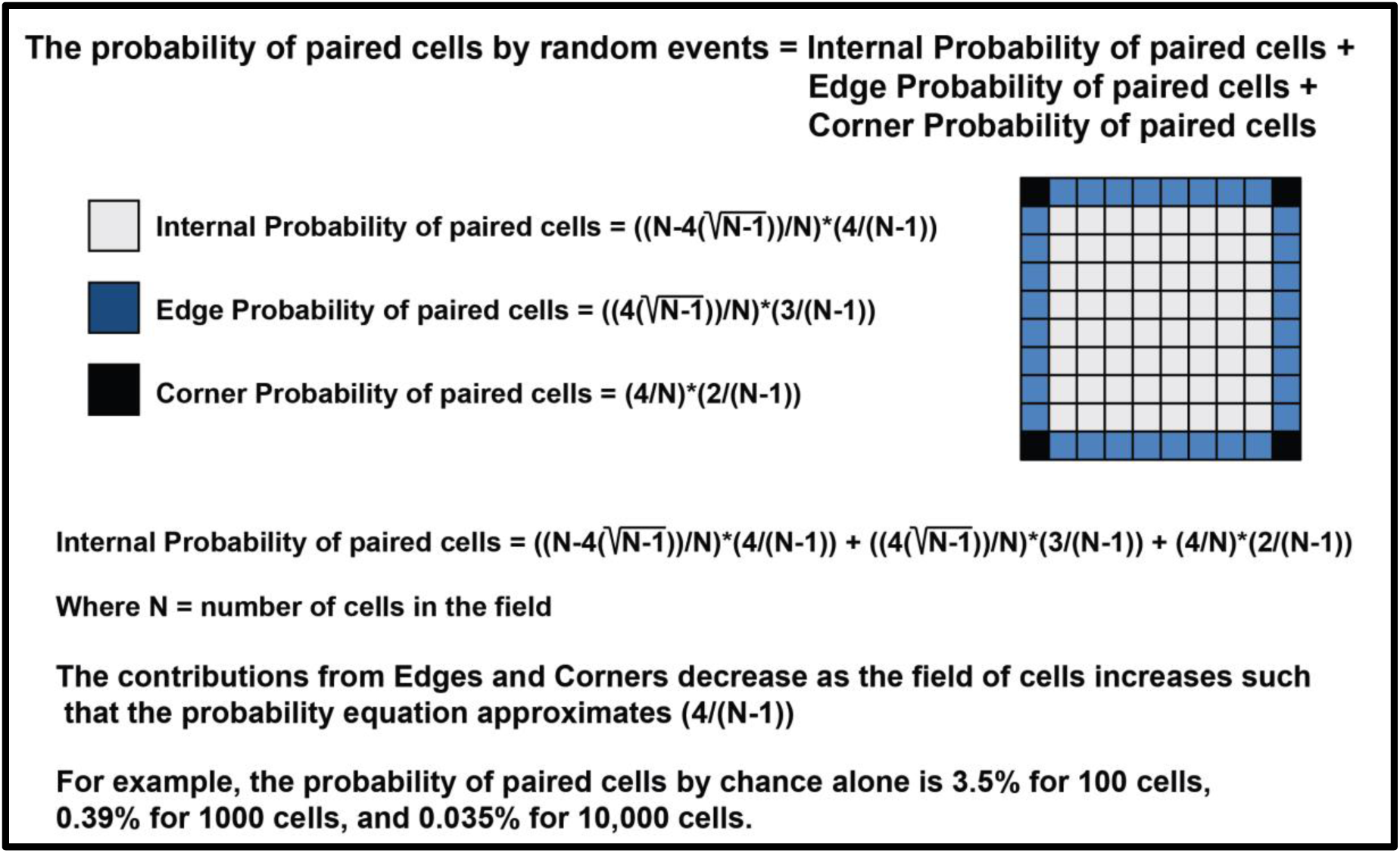
Probability estimates of paired cells occurring by chance alone. The probability equation assumes a square field of uniformly sized square cells where a pair is defined as two cells sharing one of four sides.

**Supplemental Figure 11.**
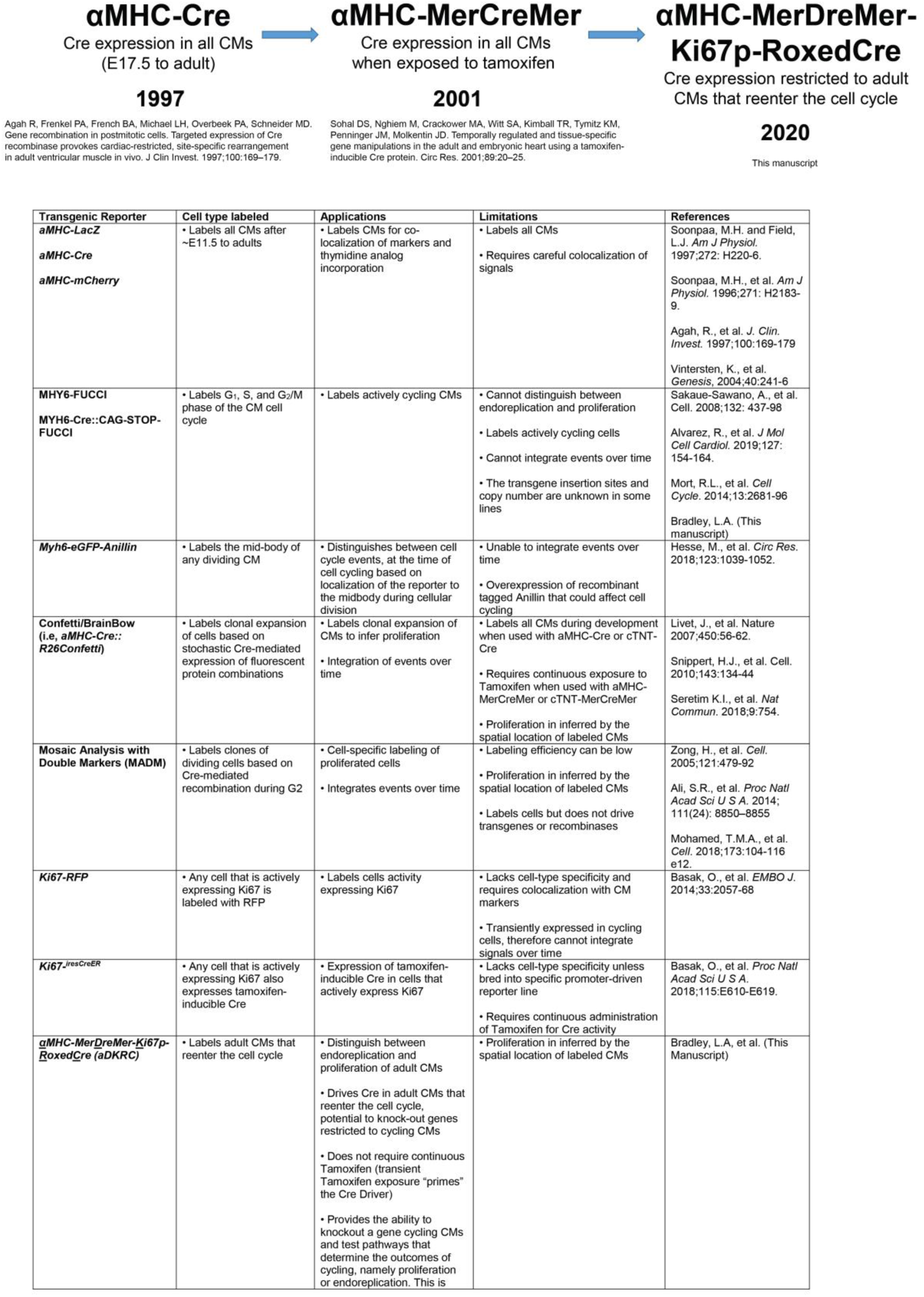
Evolution of CM-specific Cre drivers using αMHC promoters.

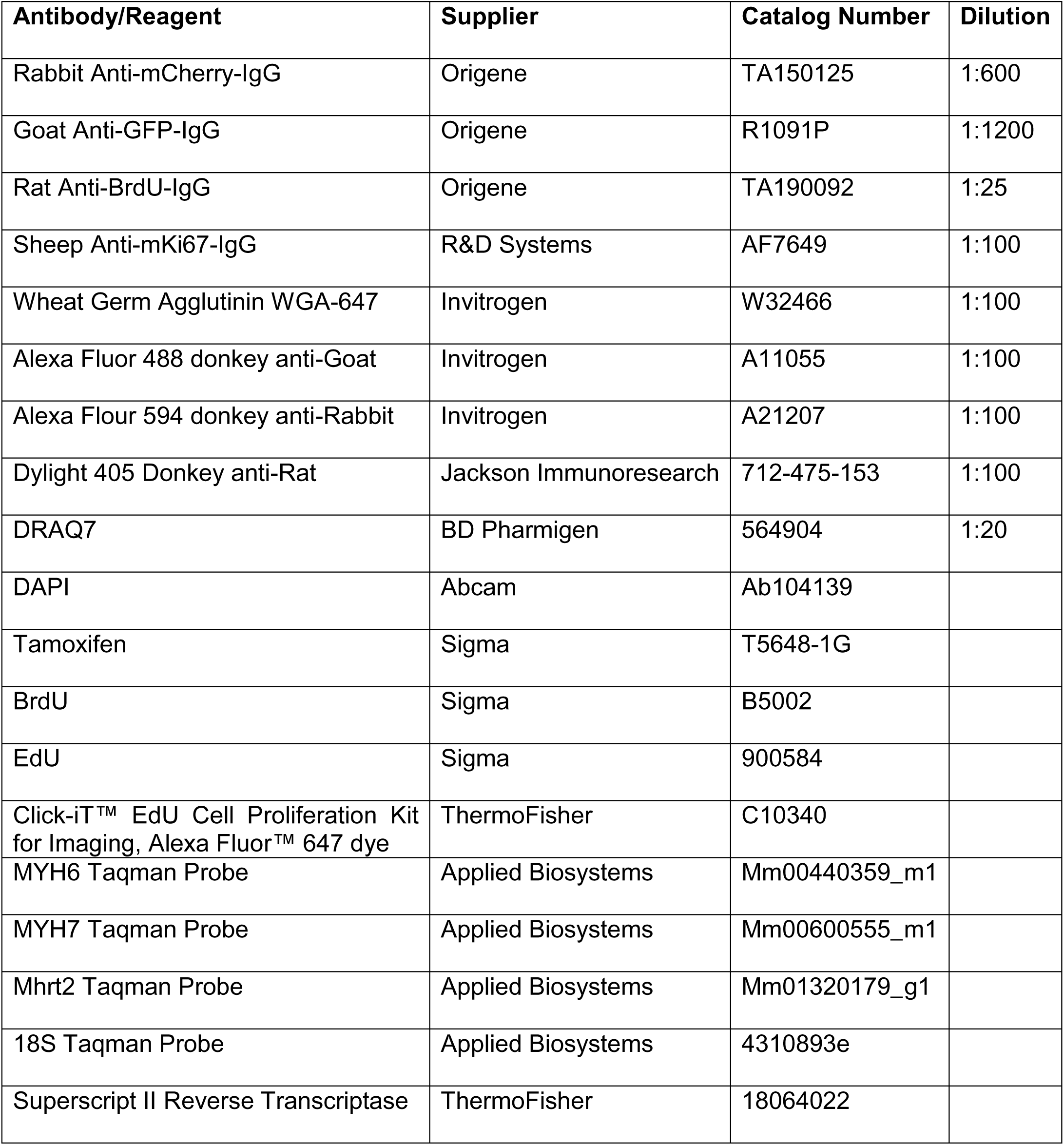
Table of Antibodies, suppliers, Catalog numbers, and Dilutions

**Table of Antibodies, suppliers, Catalog numbers, and Dilutions**
| **Antibody/Reagent** | **Supplier** | **Catalog Number** | **Dilution** |
| --- | --- | --- | --- |
| Rabbit Anti-mCherry-IgG | Origene | TA150125 | 1:600 |
| Goat Anti-GFP-IgG | Origene | R1091P | 1:1200 |
| Rat Anti-BrdU-IgG | Origene | TA190092 | 1:25 |
| Sheep Anti-mKi67-IgG | R&D Systems | AF7649 | 1:100 |
| Wheat Germ Agglutinin WGA-647 | Invitrogen | W32466 | 1:100 |
| Alexa Fluor 488 donkey anti-Goat | Invitrogen | A11055 | 1:100 |
| Alexa Flour 594 donkey anti-Rabbit | Invitrogen | A21207 | 1:100 |
| Dylight 405 Donkey anti-Rat | Jackson ImmunoResearch | 712-475-153 | 1:100 |
| DRAQ7 | BD Pharmigen | 564904 | 1:20 |
| DAPI | Abcam | Ab104139 |  |
| Tamoxifen | Sigma | T5648-1G |  |
| BrdU | Sigma | B5002 |  |
| EdU | Sigma | 900584 |  |
| Click-iT™ EdU Cell Proliferation Kit for Imaging, Alexa Fluor™ 647 dye | ThermoFisher | C10340 |  |
| MYH6 Taqman Probe | Applied Biosystems | Mm00440359_m1 |  |
| MYH7 Taqman Probe | Applied Biosystems | Mm00600555_m1 |  |
| Mhrt2 Taqman Probe | Applied Biosystems | Mm01320179_g1 |  |
| 18S Taqman Probe | Applied Biosystems | 4310893e |  |
| Superscript II Reverse Transcriptase | ThermoFisher | 18064022 |  |

